# An integrated population model for estimating the relative effects of natural and anthropogenic factors on a threatened population of Pacific trout

**DOI:** 10.1101/734996

**Authors:** Mark D. Scheuerell, Casey P. Ruff, Joseph H. Anderson, Eric M. Beamer

## Abstract

1. Assessing the degree to which at-risk species are regulated by density dependent versus density independent factors is often complicated by incomplete or biased information. If not addressed in an appropriate manner, errors in the data can affect estimates of population demographics, which may obfuscate the anticipated response of the population to a specific action.
2. We developed a Bayesian integrated population model that accounts explicitly for interannual variability in the number of reproducing adults and their age structure, harvest, and environmental conditions. We apply the model to 41 years of data for a population of threatened steelhead trout *Oncorhynchus mykiss* using freshwater flows, ocean indices, and releases of hatchery-born conspecifics as covariates.
3. We found compelling evidence that the population is under strong density dependence, despite being well below its historical population size. In the freshwater portion of the lifecycle, we found a negative relationship between productivity (offspring per parent) and peak winter flows, and a positive relationship with summer flows. We also found a negative relationship between productivity and releases of hatchery conspecifics. In the marine portion of the lifecycle, we found a positive correlation between productivity and the North Pacific Gyre Oscillation. Furthermore, harvest rates on wild fish have been sufficiently low to ensure very little risk of overfishing.
4. *Synthesis and applications.* The evidence for density dependent population regulation, combined with the substantial loss of juvenile rearing habitat in this river basin, suggests that habitat restoration could benefit this population of at-risk steelhead. Our results also imply that hatchery programs for steelhead need to be considered carefully with respect to habitat availability and recovery goals for wild steelhead. If releases of hatchery steelhead have indeed limited the production potential of wild steelhead, there are likely significant tradeoffs between providing harvest opportunities via hatchery steelhead production, and achieving wild steelhead recovery goals.

## Introduction

Managing at-risk species requires an understanding of the degree to which population dynamics are self-regulated versus driven by external factors. However, the data used to identify potentially important density-dependent and population-environment relationships are rarely, if ever, fully comprehensive or error free. Rather, imperfect detection, misidentification, and non-exhaustive sampling all lead to a somewhat distorted view of the true state of nature. For example, when not addressed in an appropriate manner, errors in population censuses may cause underestimates of recruitment (Sanz-Aguilar *et al*. 2016) or overestimates of the strength of density dependence (Knape & de Valpine 2012). Similarly, imprecision in the estimated age composition of the population also biases the estimated strength of density dependence (Zabel & Levin 2002). In a conservation context, these erroneous conclusions may directly influence the anticipated response of a population to a specific action. Therefore, proper consideration of all sources of uncertainty in the data is necessary to design robust management strategies aimed at protecting at-risk species.

The productivity and carrying capacity of a population may also vary over time and space (Thorson *et al*. 2015), and explicit consideration of external drivers can improve estimates of population dynamics under density dependent conditions (Lebreton & Gimenez 2013). For at-risk species, these exogenous factors can be used to better understand drivers of historical population demographics and help identify possible recovery options. Incorporating covariates into population models can also improve forecasts of future dynamics, especially over shorter time horizons most relevant to natural resource management (Ward *et al*. 2014). Furthermore, accelerated global change will likely create synergistic effects that complicate efforts to make reliable long-term predictions (Schindler & Hilborn 2015). Thus, any reasonable assumptions about future responses of populations should begin with an attempt to fully account for the uncertainty in population-environment relationships based on all of the available information.

Many populations of Pacific salmon (*Oncorhynchus* spp.) throughout the northwestern United States have declined markedly since the early 1900s due to a variety of causes such as habitat alteration, hydropower development, and overharvest (Ruckelshaus *et al*. 2002). For conservation purposes, Pacific salmon species are grouped into evolutionarily significant units (ESU, Waples 1991); 28 of the 49 extant ESUs of Pacific salmon are currently listed as “threatened” or “endangered” under the U.S. Endangered Species Act. As a result, a number of life-cycle models have been developed to evaluate the possible future benefits of conservation actions such as habitat restoration (e.g., Scheuerell *et al*. 2006) and the potentially negative consequences of climate change (e.g., Zabel *et al*. 2006). However, these models were assembled by first obtaining parameter values from the literature, or estimating them from disparate data sources, and then putting all of the pieces together post hoc. Consequently, they do not reflect a comprehensive assessment of the total uncertainty in population demographics.

More recently however, researchers have turned toward integrated population models (IPMs) as a means to convey the combined uncertainty in all of the data sources, which is particularly important in a conservation context (Buhle *et al*. 2018; Zipkin & Saunders 2018). IPMs are similar to state-space models in that they have specific sub-models for 1) describing the stochastic and unobservable population dynamics; and 2) addressing the noisy, incomplete data (Schaub & Abadi 2011; Maunder & Punt 2013; Yen *et al*. 2019). Although IPMs have been widely developed and applied to mammals (e.g., Eacker *et al*. 2017; Regehr *et al*. 2018) and birds (e.g., Crawford *et al*. 2018; Saunders, Cuthbert & Zipkin 2018), there are very few examples for Pacific salmon (cf., Buhle *et al*. 2018).

Here we combine incomplete data on adult abundance, age composition, and harvest into a Bayesian IPM to answer important questions relevant to management of a threatened population of anadromous steelhead trout *Oncorhynchus mykiss* Walbaum 1792 from the Skagit River basin, which drains ~6900 km^2^ in southwestern Canada and northwestern United States. Specifically, we used 39 years of age structured abundance data (1978-2018) to quantify the degree of density dependence and the effects of a specific suite of environmental drivers on intrinsic productivity within the Skagit River steelhead population. We found that although recent population censuses are well below historical estimates, the population still operates under relatively strong density dependence. We also found that streamflow during winter and releases of hatchery-reared juvenile steelhead were negatively related to wild steelhead survival, but that survival was positively related to streamflow during summers as juveniles and sea-surface temperatures experienced as adults in the North Pacific. In light of remaining uncertainty in the factors governing the population dynamics of Skagit River steelhead, this modelling framework is an effective tool for setting near term recovery goals and evaluating population level response recovery actions.

## Materials and methods

### STUDY SPECIES AND DATA

The Skagit River system is predominantly a glacially fed system that consists of a combination of rain, snow-transitional, and snow-dominated tributaries providing approximately 48 km^2^ of potential habitat suitable for spawning and rearing by wild winter run steelhead (Hard *et al*. 2015). Adult steelhead trout in the Skagit River generally enter freshwater in November through April and typically spawn in March through June. The majority of juveniles rear in freshwater for 2 years prior to migrating to sea as smolts where they spend 2 to 6 years feeding and growing before returning to freshwater as sexually mature adults to initiate spawning (i.e., they reach sexual maturity at age three through eight; ~82% mature at age four or five). Scale samples taken from wild steelhead indicate that, on average, 9% of returning adults are repeat spawners. These fish then spend a year at sea before returning again to freshwater to spawn again.

Due to a combination of logistical constraints, only a fraction of the known spawning area was surveyed for wild spawners. Specifically, standardized index reach surveys were conducted annually in only 2 of 5 major sub-basins and 13 of 63 tributaries known to support wild steelhead production. A basin-wide estimate of wild spawners was generated annually by expanding each survey to account for estimated available habitat not surveyed. Fisheries biologists in the Skagit basin generally consider the escapement estimates to be conservative: it is more likely that escapement is underestimated than overestimated because unobserved spawning sites would serve to increase abundance. Our analyses begin with surveys in 1978 and continue through 2018.

In the model described below, we evaluate several environmental indicators of survival. Specifically, flow conditions experienced by juveniles during freshwater rearing can have strong effects on their survival to adulthood via the following mechanisms: (1) spatial contraction of habitat as a result of low summer flows and high water temperatures that coincide with the period of highest metabolic demand (e.g., Crozier *et al*. 2010), and (2) habitat displacement or direct mortality resulting from peak winter flows (e.g., Irvine 1986). Therefore, we utilized long-term flow records from a gage (#12178100) located in Newhalem Creek, a snowmelt dominated stream located in the Upper Skagit River (48.66 N, 121.246 W), and maintained by the United States Geological Survey (see Appendix S1 in Supporting Information for details). Specifically, we obtained the observed maximum of daily peak flows occurring from October through March of the first freshwater rearing year, and the minimum of low summer flows occurring from June through September of the first summer of freshwater rearing.

Because conditions experienced by salmon and steelhead during their first year at sea are thought to be critical to overall survival and growth of a given year class (Beamish & Mahnken 2001), we chose the average North Pacific Gyre Oscillation index (NPGO) from January through December as an index of conditions experienced by juvenile steelhead during their first year in the ocean. Variability in the NPGO reflects annual changes in coastal upwelling and ocean circulation patterns that correlate strongly with primary and secondary production in coastal ecosystems (Di Lorenzo *et al*. 2008). Furthermore, the NPGO has been recently identified as an important indicator of early marine survival in other Pacific salmon species (Kilduff *et al*. 2015). Because most juvenile steelhead from the Skagit River migrate to sea during the spring of their second year, we lagged the NPGO indicator by two years beyond the birth year to reflect conditions experienced during the first year at sea.

From a management standpoint, we were interested in the possible effect of hatchery-reared juvenile steelhead on the productivity of wild steelhead. The Washington Department of Fish and Wildlife operates a “segregated” steelhead hatchery program (sensu Mobrand *et al*. 2005) that uses broodstock from a non-local source intentionally bred for early spawning, with the goal of minimizing temporal reproductive overlap with wild fish and hence minimizing gene flow into the wild population. Over the time series, hatchery fish were typically reared to age-1 and released in the spring (April or May) from multiple locations in the Skagit Basin. We hypothesized that hatchery fish would have the greatest potential for conspecific ecological interactions during the time juvenile steelhead are migrating to sea because observations at a juvenile fish trap (river km 27) indicate they overlap in time and space. Therefore, we assumed that a cohort born in year *t* would interact with hatchery fish released in year *t* + 2. We used the total number of juveniles released from the hatchery within a given year as our covariate.

### INTEGRATED POPULATION MODEL

The IPM we describe here expands upon models developed by others (e.g., Su & Peterman 2012; Fleischman *et al*. 2013; Winship, O’Farrell & Mohr 2014) in that we include the effects of extrinsic drivers on population dynamics. As with other IPMs, our model comprises two major components: a process model describing the production of age-specific offspring, and observation models to account for errors in the estimates of spawning escapement and age composition. Following other, more traditional analyses of Pacific salmon population dynamics, our modeling framework also assumes no consistent bias in estimates of adult spawners or age composition of returning adults.

We begin with our process model where the number of offspring born in year *t* that survive to adulthood (*R_t_*) equals the product of a nonlinear function of the number of spawning adults (*S_t_*) and a time-varying stochastic error *ε_t_*:

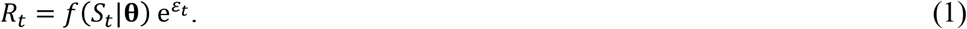

Here we consider two different forms for *f*: the Ricker model (Ricker 1954) and the Beverton-Holt model (Beverton & Holt 1957); see Fig. 1 for model forms and descriptions of their parameters and associated reference points.

**Figure 1.**
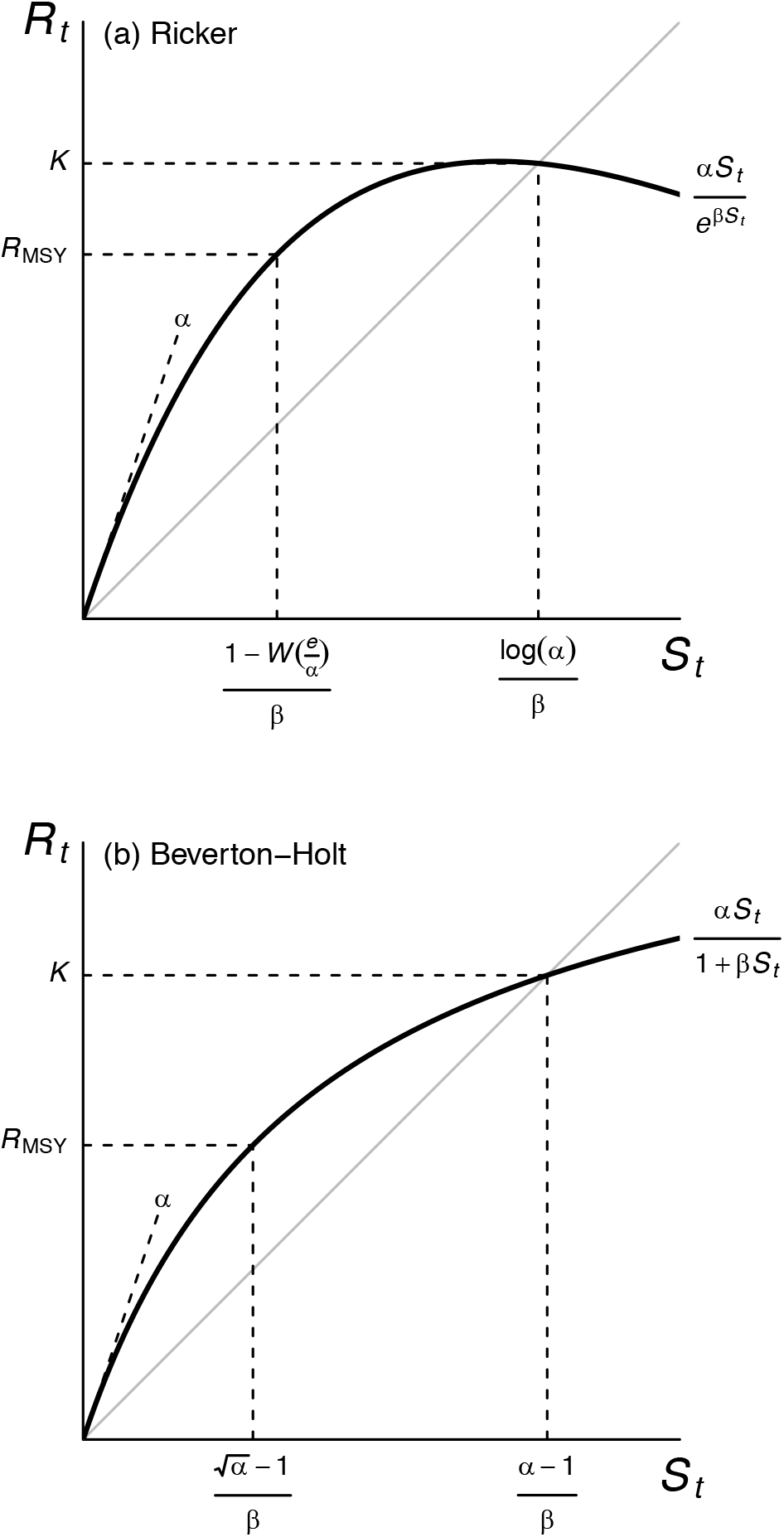
Deterministic forms of the (a) Ricker and (b) Beverton-Holt models used in the analyses (thick lines), including equations for carrying capacity (*K*) and the number of recruits corresponding to the maximum sustained yield (*R*_MSY_). The parameter *α* defines the slope at the origin, the constant *e* is Euler’s number, and *W*(·) is the Lambert function (see Scheuerell 2016 for details). The gray line is where *R_t_* = *S_t_*.

The process errors (*ε_t_*) are often assumed to be independent draws from a Gaussian distribution with a mean of zero and an unknown variance. However, the stochastic environmental drivers that the *ε_t_* are meant to represent typically show relatively strong autocorrelation over time. Thus, we compared two different distributional forms for *ε_t_* with non-zero, autocorrelated means. In the first, we assumed that

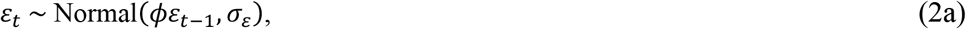

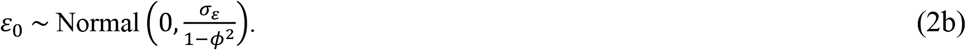

Second, we considered models where the non-zero means were also a function of the various environmental drivers important to salmon survival as discussed above. In those models,

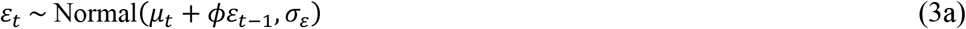

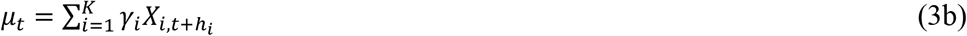

Here, *γ_i_* is the effect of covariate *X_i_* measured at time *t* and shifted by an appropriate lag *h_i_* based on the life stage that the covariate would affect most strongly. We standardized all covariates to have zero-mean and unit-variance to facilitate direct comparison of effect sizes.

The estimated numbers of fish of age *a* returning in year *t*(*N_a,t_*) is then product of the total number of brood-year recruits in year *t* − *a* from Equation (1) and the proportion of mature fish from that brood year that returned to spawn at age *a* (*π_a,t-a_*), such that

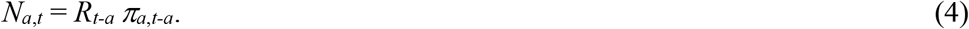

Adult steelhead from the Skagit River return as 3-8 year-olds, and therefore the vector of age-specific return rates for brood year *t* is **π**_*t*_ = [*π*_3_, *π*_4_, *π*_5_, *π*_6_, *π*_7_, *π*_8_]_*t*_, which we modeled as a hierarchical random effect whereby ***π**_t_* ~ Dirichlet(**η** *τ*). The mean vector **η** is also distributed as a Dirichlet; the precision parameter *τ* affects each of the elements in **η** such that large values of *τ* result in ***π**_t_* very close to **η** and small values of *τ* lead to much more diffuse **π**_*t*_.

The spawner-recruit models above describe a process based on the true number of spawners, but our estimates of the numbers of spawning adults necessarily contain some sampling errors due to incomplete censuses, pre-spawn mortality, etc. Therefore, we assumed that our estimates of escapement, the number of adult fish that “escape the fishery” and ultimately spawn (*E_t_*), are log-normally distributed about the true number of spawners (*S_t_*):

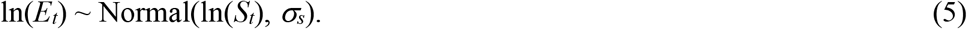

Catches of wild steelhead are closely recorded by state and tribal biologists, and so we assume the harvest is recorded without error. We then calculate *S_t_* as the difference between the estimated total run size (*N_t_*) and harvest (*H_t_*), where

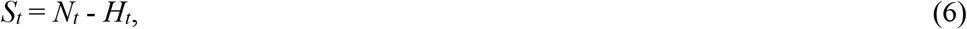

and *N_t_* is the sum of *N_a,t_* from Equation (3) over all age classes.

We obtained observations of the number of fish in each age class *a* in year *t* (*O_a,t_*) from scale analyses of 10 – 408 adults per year; no scale samples were taken in 1978-1982, 1984, and 2000. These data were assumed to arise from a multinomial process with order *Y_t_* and proportion vector **d**_*t*_, such that

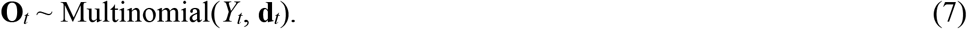

The order of the multinomial is simply the sum of the observed numbers of fish across all ages returning in year *t*:

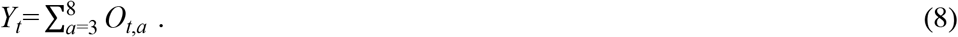

The proportion vector **d**_*t*_ for the multinomial is based on the age-specific, model-derived estimates of adult returns in year *t* (*N_a,t_*) such that

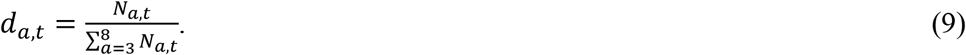

We used Bayesian inference to estimate all model parameters and the unobserved true numbers of spawners and offspring over time. We used the freely available **R** software (v3.6, R Development Core Team 2019) combined with the JAGS software (v4.2.0, Plummer 2003) to perform Gibbs sampling with 4 parallel chains of 5×10^5^ iterations. Following a burn-in period of 2.5×10^5^ iterations, we thinned each chain by keeping every 400^th^ sample to eliminate any possible autocorrelation, which resulted in 5000 samples from the posterior distributions. We assessed convergence and diagnostic statistics via the ‘CODA’ package in **R** (Plummer *et al*. 2006). Specifically, we used visual inspection of trace plots and density plots, and verified that Gelman and Rubin’s (2017) potential scale reduction factor was less than 1.1, to ensure adequate chain mixing and parameter convergence. Data support for each model was evaluated using leave-one-out cross-validation (LOO) based upon Pareto-smoothed importance sampling (Vehtari, Gelman & Gabry 2017) as implemented in the ‘loo’ package (Vehtari *et al*. 2019). All of the code and data files necessary to replicate our analyses are available in the online supporting material and at https://github.com/mdscheuerell/skagit_sthd.

## Results

We found the most data support for the Beverton-Holt form of process model, so all of the following results are based upon it (see Appendix S2 for full model selection results). Our estimates of the total population size reflect the uncertainty in the estimated numbers of adults over time, but the median values agreed quite well with the observed data (Fig. 2). As expected, the 95% credible intervals were widest in 1996 and 1997 when there were no direct estimates of spawning adults.

**Figure 2.**
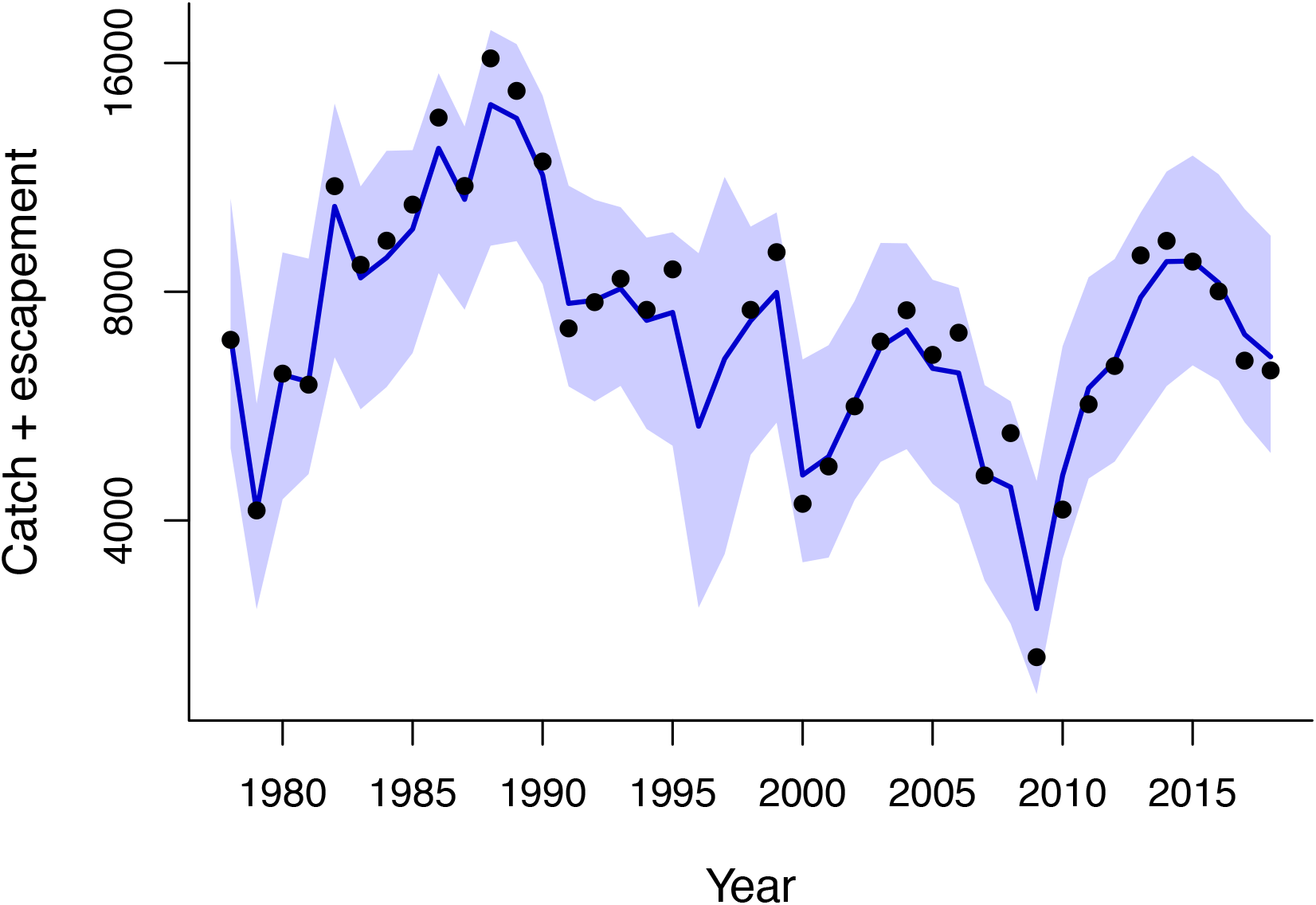
Time series of the estimated total population size (catch plus the adults that escaped to spawn). The observed data are the points; the solid line is the median estimate and the shaded region indicates the 95% credible interval.

The population dynamics of steelhead in the Skagit River are currently under density-dependent regulation, despite their numbers being well below historical censuses, and there is considerable uncertainty in the relationship between spawning adults and their surviving offspring (Fig. 3). The median of a (i.e., the slope of the relationship at the origin) was 6.8 offspring per spawner, but a lack of data at low spawner abundance led to considerable uncertainty in the estimate (Fig. 3b). The lower 95% credible interval was about 1.5 offspring per spawner, which is still above replacement, while the upper 95% credible interval was 44 offspring per parent. On the other hand, our estimates of carrying capacity (*K*) were much more precise, with a median of about 7400 adults and 95% credible interval of approximately 6100 to 10 900 adults (Fig. 3c).

**Figure 3.**
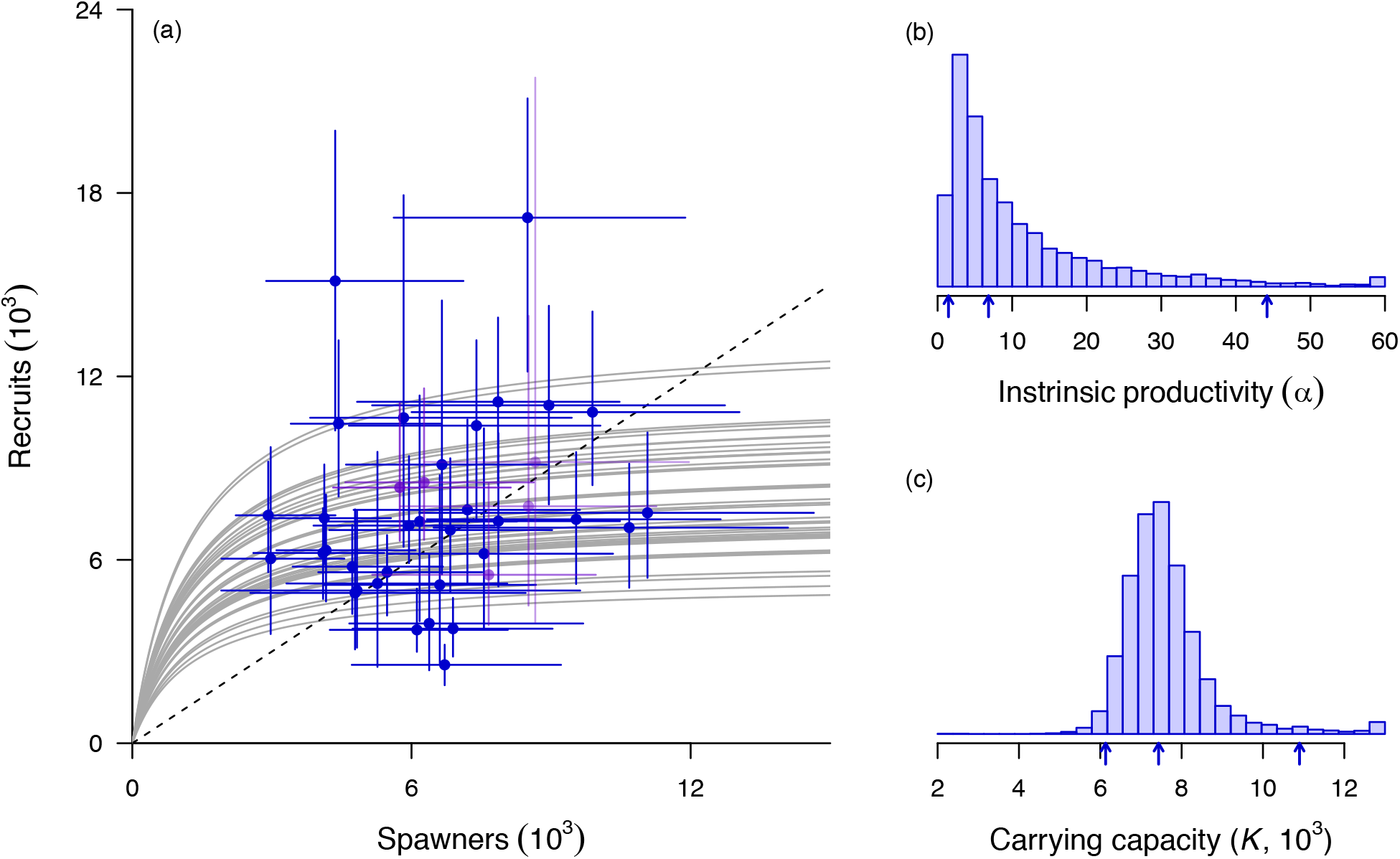
Relationship between the number of spawning adults and their subsequent surviving offspring (recruits), assuming mean values for all covariates (a); and the estimated posterior distributions for the intrinsic productivity (b) and carrying capacity (c). Points in (a) are medians of the posterior estimates; error bars indicate the 95% credible intervals. Blue points are for estimates with complete broods; purple points are for the most recent years with incomplete broods. Gray lines show the median relationship for each of the 41 years in the time series based on annual model estimates of productivity. Note that for plotting purposes only in (b) and (c), the density in the largest bin for each parameter contains counts for all values greater than or equal to it. Vertical arrows under the x-axes in (b) and (c) indicate the 2.5^th^, 50^th^, and 97.5^th^ percentiles.

There were varying effects of the three environmental covariates on population productivity (Fig. 4). Peak winter flows were negatively related to survival, suggesting high discharge events may transport juveniles downstream to lower quality habitats, or lead to direct mortality from channel avulsion or movement of sediment, wood, and other debris. The median of the posterior distribution was −0.11 (Fig. 4e), which means that a 1 SD increase in flow above the mean (i.e., from ~41 m^3^ s^-1^ to ~68 m^3^ s^-1^) would translate into a 11% decrease in offspring per parent. Conversely, the effect of low summer flows was positive (Fig. 4f), possibly indicative of greater rearing habitat (the median estimate was 0.08 with a 95% credible interval of −0.09 to 0.25). The NPGO had a similar effect to summer flow (Fig. 4g), suggesting warmer waters in the North Pacific are better for steelhead survival (median equals 0.09 with a 95% credible interval of −0.08 to 0.27.

**Figure 4.**
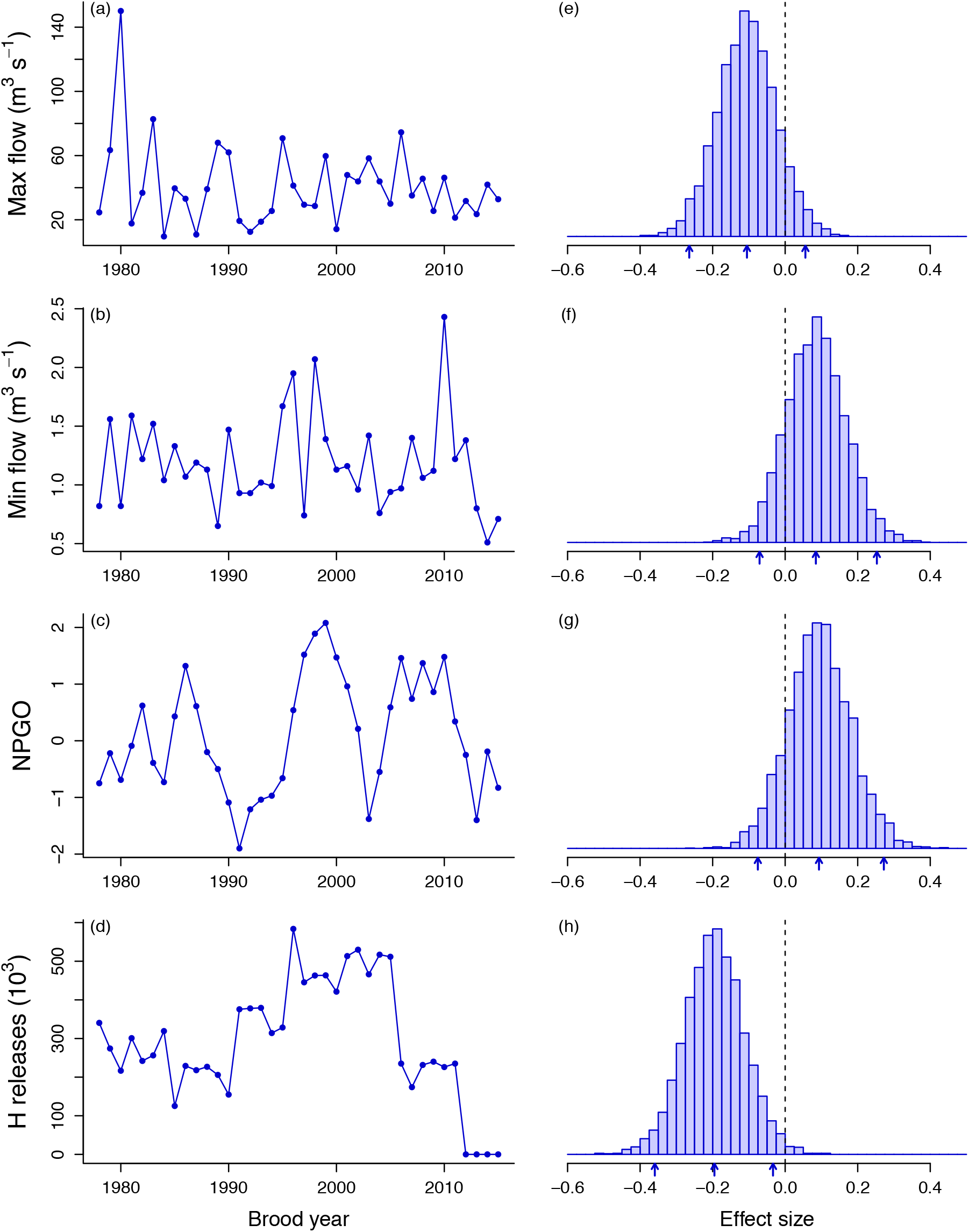
Time series of the environmental covariates used in the model (a-d), and their standardized effects on population productivity (e-g). Small arrows under histograms denote the 2.5^th^, 50^th^, and 97.5^th^ percentiles of the posterior distribution.

We also found that the number of hatchery juveniles released into the river during the time that wild juveniles were migrating to sea was negatively related to productivity (Fig. 4h). The median effect size was −0.20, which means that a 1 SD increase in the number of hatchery juveniles released (i.e., from 328 000 to 452 000 fish) would, on average, result in a 18% decrease in survival to adulthood. Notably, hatchery production experienced three distinct phases over time (Fig. 4d): a low period between brood year 1978 and 1990 (range = 125 000 to 340 000 smolts), an increasing and high period between 1991 and 2005 (range = 314 000 to 584 000), and a decreasing period beginning in 2006 (range = 0 to 240 000 smolts).

The remaining, unexplained environmental variance was indeed highly autocorrelated over time (Fig. 5). The process residuals were generally positive during the late 1970s and early 1980s when the population was growing (Fig. 2), they were near zero during the stable period of the 1990s, and then largely negative as the population primarily declined through the 2000s.

**Figure 5.**
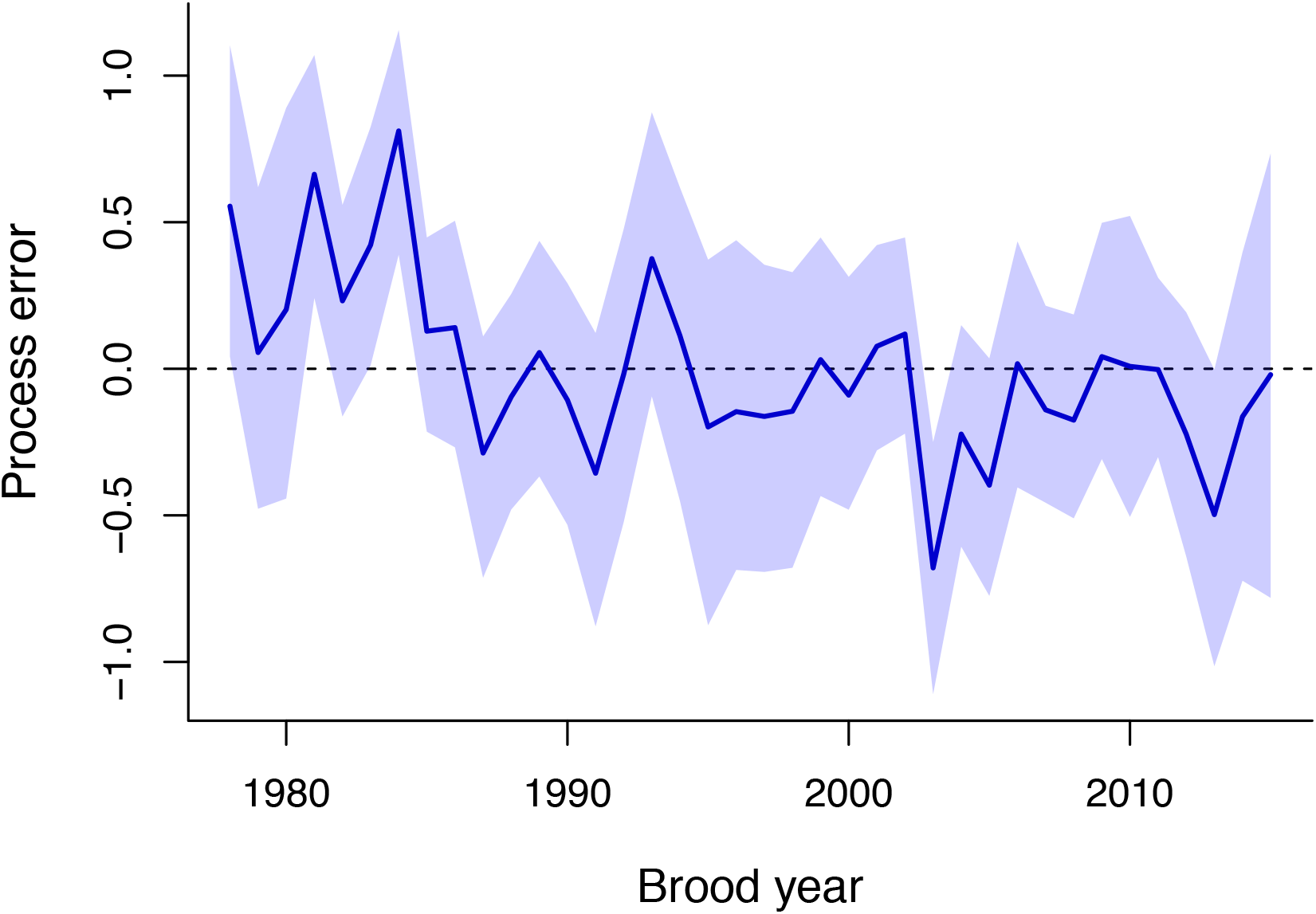
Time series of the estimated process errors, which represent the population’s productivity after accounting for the effects of density dependence and environmental covariates. The solid line is the median estimate and the shaded region indicates the 95% credible interval.

Based on our estimates of biological reference points, Skagit River steelhead appear to be managed along a rather conservative harvest management perspective. The optimal yield profiles suggest it would take approximately 2000 to 3000 spawning adults to produce the maximum sustainable yield (Fig. 6a), but very few years have ever fallen below that throughout the time period presented here (i.e., the average number of spawning adults has been two to three times greater). In other words, the realized harvest rates have been kept low enough to insure very little risk of overfishing (Fig. 6b).

**Figure 6.**
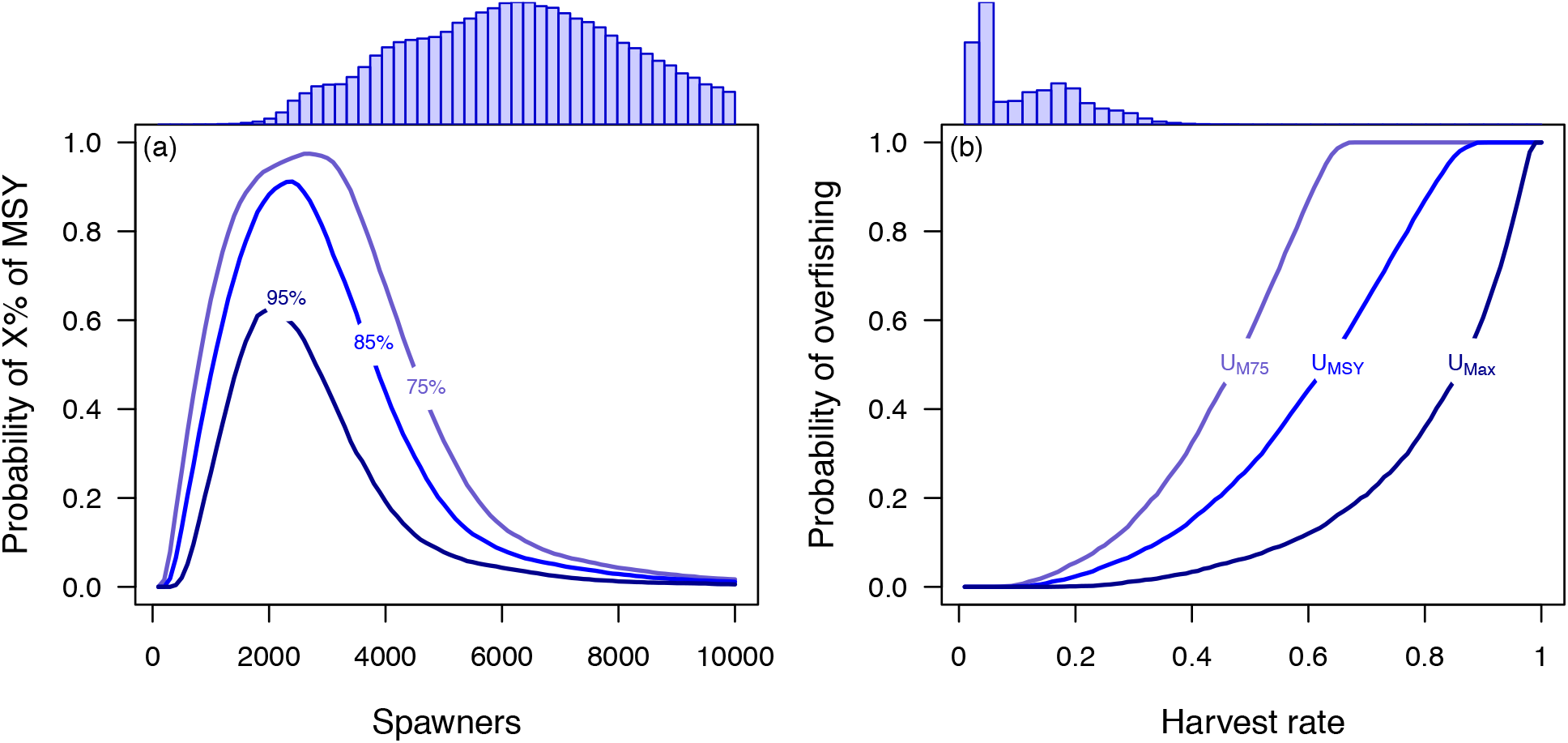
Plots of (a) the probability that a given number of spawners produces average yields achieving 95%, 85%, or 75% of the estimated maximum sustainable yield (MSY); and (b) the cumulative probability of overfishing the population, based on harvest rates equal to those at 75% of MSY, at MSY, and at the maximum per recruit. The histograms above (a) and (b) are distributions of the posterior estimates for the number of spawners and harvest rates, respectively; the histogram in (a) has been truncated at 10^4^.

## Discussion

In territorial species such as steelhead trout, competition for limited resources commonly results in density dependent growth and survival amongst juveniles (Imre, Grant & Keeley 2004). Our analysis suggests that such effects have scaled up to the entire population level to govern patterns of steelhead productivity in the Skagit River basin. Importantly, we found strong evidence for density dependent interactions despite the fact that contemporary population censuses are well below historical estimates (Gayeski, McMillan & Trotter 2011). Similar results have been observed in populations of coho salmon *Oncorhynchus kisutch* Walbaum 1792 in Oregon (Buhle *et al*. 2009) and in populations of Chinook salmon *Oncorhynchus tshawytscha* Walbaum 1792 in Idaho (Thorson *et al*. 2013). Although we cannot be certain of the exact life-stage at which density dependent processes occurred, the freshwater juvenile stage seems likely given the extended duration of freshwater rearing typical for this species. When steelhead populations reach low numbers, the spatial contraction of spawners may exacerbate the effects of density dependence because their newly emerged offspring do not have the mobility to access other vacant habitats (Atlas *et al*. 2015). The evidence for density dependence presented here, combined with the substantial loss of juvenile rearing habitat in the Skagit River basin (Beechie, Beamer & Wasserman 1994), suggests that habitat restoration efforts, such as reconnecting floodplain habitats and improving riparian functioning (Beechie, Pess & Roni 2008), may benefit this population of steelhead.

Fluctuating environments can also affect population dynamics through density independent mechanisms, and anadromous salmon must contend with many different and unpredictable habitats over their lifespan. Our results indicate that in the freshwater environment, large flow events during winter negatively affect steelhead productivity. Unfortunately, this may portend an uncertain future for these fish. In a recent study, Lee et al. (2015) estimated that future climate change in the Skagit River basin would create increased winter flows. These changes in hydrology will likely result in much greater exposure of steelhead to extreme high flow events due to their duration, intensity, and timing (Wade *et al*. 2013). Other evidence already exists that freshwater discharge from Puget Sound rivers has become much more variable, with notable negative effects on Chinook salmon *Oncorhynchus tshawytscha* Walbaum 1792 (Ward *et al*. 2015). Furthermore, although we found a somewhat weaker relationship between low summer flow and productivity, extreme low-flow events are projected to occur at a higher frequency in the future (Lee *et al*. 2015).

We found evidence of positive effects of NPGO on survival, which comports with previous studies that have made rather compelling cases for a strong positive relationship between the NPGO and salmon survival (Kilduff *et al*. 2015). The NPGO is a synoptic measure of ocean conditions over a large region of the North Pacific Ocean (Kilduff *et al*. 2015), so we cannot say where and when, exactly, the effects of the ocean environment most manifest themselves. Recent evidence also indicates that steelhead smolts suffer high mortality during their relatively brief migration through Puget Sound (Moore *et al*. 2015), possibly due to predation by marine mammals (Berejikian, Moore & Jeffries 2016). Notably, too, the residual process errors not captured by our covariates (*w_t_* in Equation 1) were correlated with the estimated marine survival of Skagit River hatchery steelhead (median Pearson’s correlation coefficient = 0.29; 95% credible interval = [0.03, 0.50]), suggesting marine processes not captured by our covariates likely influenced productivity.

Among the various mitigation measures to address salmon declines, artificial propagation of salmon has been used widely for more than a century. Nevertheless, research in other river systems points to negative ecological effects of hatchery fish on wild Pacific salmon, including populations coho salmon (Buhle *et al*. 2009), and Chinook salmon (Levin, Zabel & Williams 2001). Our results provide further evidence that large releases of hatchery-reared juvenile steelhead have had a negative effect on productivity of wild steelhead, although we note some researchers have used an approach similar to ours and found no hatchery effect on productivity (Courter *et al*. 2019; Nelson *et al*. 2019). Our study was unable determine the mechanism responsible for the correlation between hatchery releases and wild steelhead productivity. In fact, very few empirical studies have been conducted at the appropriate spatial and temporal scales necessary to directly quantify the hypothesized mechanisms by which negative ecological interactions between hatchery and wild fish may occur (Weber & Fausch 2003). That said, however, competition for limiting freshwater food and habitat resources (Berejikian *et al*. 2000) is a plausible mechanism, either during the relatively brief period of overlap during downstream migration (ca. 2 – 4 weeks), or a more prolonged effect of any hatchery fish that do not migrate to sea, but instead “residualize” within freshwater. Additionally, predators are known to respond numerically to their prey, and it is possible that large numbers of hatchery fish attracted additional predators (Kostow 2009). Although breeding by hatchery individuals that stray onto natural spawning grounds may reduce the fitness of a wild population via gene flow from the hatchery stock into the wild population (Araki, Cooper & Blouin 2009), our study only considered within-cohort effects. Thus, it seems unlikely that a trans-generational genetic effect was the mechanism for the observed negative association between hatchery releases and wild productivity.

Throughout the Puget Sound region, steelhead have been exposed to varying degrees of influence by hatchery fish over the past 100 years, but they share the marine rearing environment, and thus have experienced relatively similar ocean conditions during the same time period. The marked decreases in abundance observed in many of these populations from the late 1980s to the late 2000’s, including the Skagit, mirrors observations of a general declining trend in marine survival of hatchery conspecifics across the same time period, suggesting some larger, unmeasured forces have been at work (Kendall, Marstrom & Klungle 2017). Furthermore, in response to the declining abundance of wild Skagit River steelhead coupled with declining marine survival of hatchery steelhead, fisheries managers increased hatchery production to replace lost fishing opportunities. Thus, it is plausible that declining wild productivity was simply coincident with higher hatchery production, rather than a consequence of it. It is also possible that multicollinearity among measured and unmeasured covariates increased the estimated effect sizes.

The life history complexity of steelhead may not lend well to the use of traditional spawner recruit models such as the forms used in this study. Notably, steelhead exhibit significant phenotypic plasticity with respect to adopting partial migration strategies, with unknown proportions of a given cohort adopting a non-anadromous resident life history type (Kendall *et al*. 2015). Given that only anadromous individuals are included in the annual derivation of age structured abundance, there may be a large component of each cohort that is missed which likely resulted in substantial observation error not captured in our models. Therefore, caution should be used when interpreting the spawner recruit relationships and resulting management reference points presented here. Furthermore, future research should aim at quantifying the contribution of individuals adopting the resident life history type to overall productivity. Without these estimates, accurate assessments of the status of steelhead populations may not be possible. The “precautionary approach” to fisheries management aims to balance the trade-off between catch and the risk of over-fishing such that minimizing the risk of overfishing takes precedence (Hilborn *et al*. 2001). Our Bayesian state-space model provides a formal means for estimating the probability of fishing in a sustainable manner. We found compelling evidence that harvest rates for wild steelhead in the Skagit River basin over the time period considered here have been well below those that would drive the population toward extinction. This result, combined with the strong indication of density dependence, lends further support to the notion that habitat improvements may benefit this population most. However, some caution is warranted because we may have overestimated the biological reference points by not fully accounting for repeat spawners.

Here we have demonstrated how to use incomplete information about the abundance and age structure of a population to estimate density dependent population dynamics in light of natural and human-induced variability in the environment. Our study adds to the growing body of evidence that habitat, hatchery practices, and environmental variability are intricately linked in affecting productivity of wild Pacific salmon stocks. Future research should focus on quantifying habitat limitation on productivity at specific life stages to better focus restoration actions needed to recover wild steelhead. Our modeling framework also allowed us to assess the degree to which hatchery and harvest management actions are likely to affect the long-term viability of the population. Our results suggest that hatchery program goals for steelhead need to be considered carefully with respect to recovery goals and the quantity and quality of steelhead habitat. If releases of non-local origin hatchery steelhead have indeed limited the production potential of wild steelhead, there are likely significant tradeoffs between providing harvest opportunities via hatchery steelhead production and achieving wild steelhead recovery goals.

## Supporting information

Appendix 1 - Data retrieval

Appendix 2 - Model fitting

Appendix 3 - Summarize results

## Acknowledgements

We thank Eric Buhle and Jim Thorson for helpful discussions about model development. We also thank Rebecca Bernard, Pete Kairis, Brett Barkdull, Andrew Fowler, and numerous WDFW, Sauk-Suiattle, Swinomish, and Upper Skagit tribal biologists who compiled the escapement and age data. Annette Hoffman, Dan Rawding, Kris Ryding, and James Scott provided constructive criticism. This research was funded in part by a NOAA Fisheries And The Environment (FATE) grant (#15-01), and an Environmental Protection Agency Tribal Capacity grant (#PA-00J320-01).

## Authors’ Contributions

MS, CR and JA conceived the ideas and designed methodology; MS and CR analysed the data; MS and CR led the writing of the manuscript. All authors contributed critically to the drafts and gave final approval for publication.

## Data Accessibility

All of the fish data have been archived at Figshare and are available via the following links:

abundance (https://dx.doi.org/10.6084/m9.figshare.3458183.v1);
age composition (https://dx.doi.org/10.6084/m9.figshare.3458204.v1);
harvest (https://dx.doi.org/10.6084/m9.figshare.3458189.v1); and
hatchery releases (https://dx.doi.org/10.6084/m9.figshare.3457163.v1).

The river flow data are available from the United States Geological Survey National Water Information System (http://waterdata.usgs.gov/nwis). The North Pacific Gyre Oscillation data are available from Emanuele Di Lorenzo at Georgia Technical University (http://www.o3d.org/npgo/).

## SUPPORTING INFORMATION

Additional Supporting Information may be found in the online version of this article:

**Appendix S1**. Instructions for retrieving and archiving the environmental covariates.

**Appendix S2**. Model definitions, model fitting, and model evaluation.

**Appendix S3**. Steps to recreate figures from main text.

