## Appendix 1 - Data retrieval for "An integrated population model for estimating the relative effects of natural and anthropogenic factors on a threatened population of Pacific trout"

### Appendix S1. Instructions for retrieving and archiving the environmental covariates.

Supporting information for Scheuerell et al.

**4 Archive covariates 6**

This is version 0.19.07.08.

#### 1 Background

This appendix describes how to retrieve the environmental covariates used in our analyses. After reading in the raw data, summarizing them (if necessary), and trimming them to the appropriate time frame, they table of covariates is written to a `.csv` file.

All of the analyses require the R software (v3.4.3 or later) for data retrieval and processing. We also need the **readr** and **here** packages, which are not included with the base installation of **R**.

```
if(!require("readr")) {
  install.packages("readr")
  library("readr")
}
if(!require("here")) {
  install.packages("here")
  library("here")
}
## set data dir
datadir <- here("data")
```

#### 2 User inputs

We begin by supplying values for the following parameters, which we use for trimming and lagging the covariates to the appropriate years.

```r
## first & last years of fish data
yr_frst <- 1978
yr_last <- 2018

## min & max adult ages (years)
age_min <- 3
age_max <- 8

## time lags (years) for covariates
flow_lag <- 1
marine_lag <- 2
hrel_lag <- 2

## number of years for run forecasts
n_fore <- 0
```

### 3 Retrieve covariates

Our analysis investigates 4 covariates as possible drivers of the population's instrinic growth rate:

1. Maximum river discharge in winter;
2. Minimum river discharge in summer;
3. North Pacific Gyre Oscillation;
4. Releases of hatchery-born juveniles.

#### 3.1 River discharge

We begin by getting the daily flow data from the US Geological Service National Water Information System. We will use the direct link to the gage data from the Skagit River near Mount Vernon, WA (#12178100), beginning with the first year of fish data.

```r
## flow gage ID
flow_site <- 12178100
## get URL for flow data from USGS
flow_url <- paste0("https://waterdata.usgs.gov/nwis/dv",
                   "?cb_00060=on",
                   "&format=rdb",
                   "&site_no=",flow_site,
                   "&begin_date=",yr_frst,"-01-01",
                   "&end_date=",yr_last,"-12-31")
```

Next we retrieve the raw data file and print its metadata.

```r
## raw flow data from USGS
flow_raw <- read_lines(flow_url)
## lines with metadata
hdr_flow <- which(lapply(flow_raw, grep, pattern = "\\#")==1, arr.ind = TRUE)
## print flow metadata
print(flow_raw[hdr_flow], quote = FALSE)
```

```
##  [1] # ---------------------------------- WARNING ----------------------------------------
##  [2] # Some of the data that you have obtained from this U.S. Geological Survey database
##  [3] # may not have received Director's approval. Any such data values are qualified
##  [4] # as provisional and are subject to revision. Provisional data are released on the
##  [5] # condition that neither the USGS nor the United States Government may be held liable
##  [6] # for any damages resulting from its use.
##  [7] #
##  [8] # Additional info: https://help.waterdata.usgs.gov/policies/provisional-data-statement
##  [9] #
## [10] # File-format description:  https://help.waterdata.usgs.gov/faq/about-tab-delimited-ou
## [11] # Automated-retrieval info: https://help.waterdata.usgs.gov/faq/automated-retrievals
## [12] #
## [13] # Contact:  
## [14] # retrieved: 2018-11-28 12:21:31 EST       (vaww02)
## [15] #
## [16] # Data for the following 1 site(s) are contained in this file
## [17] #    USGS 12178100 NEWHALEM CREEK NEAR NEWHALEM, WA
## [18] # -----------------------------------------------------------------------------------
## [19] #
## [20] # Data provided for site 12178100
## [21] #            TS   parameter     statistic     Description
## [22] #        149343       00060     00003     Discharge, cubic feet per second (Mean)
## [23] #
## [24] # Data-value qualification codes included in this output:
## [25] #     A  Approved for publication -- Processing and review completed.
## [26] #     P  Provisional data subject to revision.
## [27] #     e  Value has been estimated.
## [28] #
```

Lastly, we extract the actual flow data for the years of interest and inspect the file contents.

```
## flow data for years of interest
dat_flow <-  read_tsv(flow_url,
                      col_names = FALSE,
                      col_types = "ciDdc",
                      skip = max(hdr_flow)+2)
colnames(dat_flow) <- unlist(strsplit(tolower(flow_raw[max(hdr_flow)+1]),
                                      split = "\\s+"))
head(dat_flow)
```

```
## # A tibble: 6 x 5
##   agency_cd  site_no datetime   `149343_00060_00003` `149343_00060_00003_cd`
##   <chr>        <int> <date>                    <dbl> <chr>
## 1 USGS      12178100 1978-01-01                   58 A
## 2 USGS      12178100 1978-01-02                   55 A
## 3 USGS      12178100 1978-01-03                   56 A
## 4 USGS      12178100 1978-01-04                   66 A
## 5 USGS      12178100 1978-01-05                   90 A
## 6 USGS      12178100 1978-01-06                   87 A
```

We only need the 3rd and 4th columns, which contain the date (`datetime`) and daily flow measurements (`149343_00060_00003`). We will rename them to `date` and `flow`, respectively, and convert the flow units from "cubic feet per second" to "cubic meters per second".

```
## keep only relevant columns
dat_flow <- dat_flow[c("datetime", grep("[0-9]$", colnames(dat_flow), value = TRUE))]
## nicer column names
colnames(dat_flow) <- c("date","flow")
## convert cubic feet to cubic meters
dat_flow$flow <- dat_flow$flow / 35.3147
## flow by year & month
dat_flow$year <- as.integer(format(dat_flow$date,"%Y"))
dat_flow$month <- as.integer(format(dat_flow$date,"%m"))
dat_flow <- dat_flow[,c("year","month","flow")]
```

##### 3.1.1 Winter maximum

We are interested in the maximum of the daily peak flows from October through March during the first year that juveniles are rearing in streams. This means we need to combine flow values from the fall of year $t$ with those in the winter and spring of year $t + 1$. We also need to shift the flow data forward by 1 year so they align with the juvenile life stage. Therefore, the flow time series will begin in 1978 and end in 2016.

```
## autumn flows in year t
flow_aut <- subset(dat_flow, (month>=10 & month<=12)
                   & year >= yr_frst & year <= yr_last-age_min+n_fore)
## spring flows in year t+1
flow_spr <- subset(dat_flow,
                   (month>=1 & month<=3)
                   & year >= yr_frst+flow_lag
                   & year <= yr_last-age_min+n_fore+flow_lag)
## change spr year index to match aut
flow_spr[,"year"] <- flow_spr[,"year"] - flow_lag
## combined flows indexed to brood year & calculate max flow
dat_flow_wtr <- aggregate(flow ~ year, data = rbind(flow_aut,flow_spr), max)
dat_flow_wtr[,"flow"] <- round(dat_flow_wtr[,"flow"], 1)
## change year index to brood year
dat_flow_wtr[,"year"] <- dat_flow_wtr[,"year"]
## for plotting purpose later
colnames(dat_flow_wtr)[2] <- "flow_wtr"
```

##### 3.1.2 Summer minimum

Retrieving the minimum flow juveniles would experience during their first summer (June through September) is straightforward.

```
## summer flows in year t
flow_sum <- subset(dat_flow, (month>=6 & month<=9)
                   & year >= yr_frst+flow_lag
                   & year <= yr_last-age_min+n_fore+flow_lag)
```

```r
## change year index to brood year
flow_sum[,"year"] <- flow_sum[,"year"] - flow_lag
## combined flows indexed to brood year & calculate min flow
dat_flow_sum <- aggregate(flow ~ year, data = flow_sum, min)
dat_flow_sum <- round(dat_flow_sum, 2)
## for plotting purpose later
colnames(dat_flow_sum)[2] <- "flow_sum"
```

#### 3.2 North Pacific Gyre Oscillation

We used the monthly NPGO data provided by Emanuele Di Lorenzo of the Georgia Institute of Technology, which are available here. We begin by downloading the raw NPGO data and viewing the metadata.

```r
## URL for NPGO data
url_NPGO <- "http://www.o3d.org/npgo/npgo.php"
## raw NPGO data
NPGO_raw <- read_lines(url_NPGO)
## line with data headers
hdr_NPGO <- which(lapply(NPGO_raw,grep,pattern="YEAR")==1, arr.ind = TRUE)
## print PDO metadata
print(NPGO_raw[seq(hdr_NPGO)],quote = FALSE)
```

```
##  [1]
##  [2] <html>
##  [3] <body>
##  [4]
##  [5] <pre>#  Last update 23-Feb-2018 by E. Di Lorenzo
##  [6] #  NPGO index monthly averages
##  [7] #  from Jan-1950  to  Feb-2018
##  [8] #
##  [9] #  WARNING: Values after Dec-2004 are updated
## [10] #  using Satellite SSHa from AVISO Delayed Time product.
## [11] #  http://opendap.aviso.oceanobs.com/thredds/dodsC/dataset-duacs-dt-global-allsat-msla
## [12] #
## [13] #  PRELIMINARY: Values after May-2017 are preliminary and updated
## [14] #  using Satellite SSHa from AVISO Near Real Time product.
## [15] #  http://opendap.aviso.oceanobs.com/thredds/dodsC/dataset-duacs-nrt-over30d-global-al
## [16] #
## [17] #  The update is performed by taking the NPGO spatial pattern of Di Lorenzo et al. 200
## [18] #  computed over the period 1950-2004, and projecting the AVISO Satellite SSHa.
## [19] #  During the pre-processing of the AVISO data, we remove the seasonal cycle based on
## [20] #  the 1993-2004 seasonal means.
## [21] #
## [22] #  AVISO PRODUCT UPDATE Summer 2014: AVISO has released a re-processed dataset for the
## [23] #  Starting from the November 2014, the NPGO index is computed with this updated datas
## [24] #  values from 2004 onward have been recomputed with very minor differences from previ
## [25] #
## [26] #  Ref:
```

```
## [27] #  Di Lorenzo et al., 2008: North Pacific Gyre Oscillation
## [28] #  links ocean climate and ecosystem change, GRL.
## [29] #
## [30] #      YEAR            MONTH        NPGO index
```

Next, we extract the actual NPGO indices for the years of interest and inspect the file contents. We also want the average NPGO annual index from January 1 through December 31 during the first year that the juvenile steelhead are in the ocean (i.e., during their second year of life). Therefore, we need NPGO values from `yr_frst + marine_lag == 1980` through `yr_last - age_min + n_fore + marine_lag == 2017`.

```
## number of years of data
n_yrs <- yr_last - yr_frst + 1
## NPGO data for years of interest
dat_NPGO <- read_table(url_NPGO, col_names = FALSE,
                       skip = hdr_NPGO + (yr_frst-1950)*12,
                       n_max = (n_yrs-1)*12)
colnames(dat_NPGO) <- c("year","month","NPGO")
## select only years of interest indexed by brood year
dat_NPGO <- dat_NPGO[dat_NPGO$year >= yr_frst+marine_lag &
                     dat_NPGO$year <= yr_last-age_min+n_fore+marine_lag,]
dat_NPGO <- aggregate(dat_NPGO$NPGO, by = list(year = dat_NPGO$year), mean)
dat_NPGO <- data.frame(year = seq(yr_frst,yr_last-age_min+n_fore),
                       NPGO = dat_NPGO[,2])
dat_NPGO[,"NPGO"] <- round(dat_NPGO[,"NPGO"], 2)
```

##### 3.3 Hatchery releases

The numbers of hatchery fish released each year are provided in `/data/skagit_sthd_hrel.csv`. They have already been lagged 2 years (i.e., brood year + 2) to account for the potential competitive interactions during their juvenile life stage.

```
## get releases
dat_hrel <- read_csv(file.path(datadir, "skagit_sthd_hrel.csv"))
dat_hrel <- subset(dat_hrel, dat_hrel$year <= max(dat_NPGO$year))
```

#### 4 Archive covariates

The last thing we will do is combine the covariates into one data frame and write them to a file for use in the analysis.

```
## combine covariates
dat_cvrs <- Reduce(function(...) merge(..., all = TRUE),
                   list(dat_flow_wtr,
                        dat_flow_sum,
                        dat_NPGO,
                        dat_hrel))
## check table of covariates
head(dat_cvrs)
```

```
##   year flow_wtr flow_sum  NPGO   Hrel
## 1 1978     24.6     0.82 -0.75 340416
## 2 1979     63.4     1.56 -0.22 274090
## 3 1980    150.1     0.82 -0.69 216537
## 4 1981     17.7     1.59 -0.09 300942
## 5 1982     36.8     1.22  0.62 241931
## 6 1983     82.7     1.52 -0.39 256238
```

```
## write covariates to a file
write_csv(dat_cvrs, file.path(datadir, "skagit_sthd_covars.csv"))
```
