## Appendix 2 - Model fitting for "An integrated population model for estimating the relative effects of natural and anthropogenic factors on a threatened population of Pacific trout"

### Appendix S2. Model definitions, model fitting, and model evaluation.

Supporting information for Scheuerell et al.

#### Contents


**1 Background** **1**

**2 User inputs** **3**

**3 Loading the fish data** **3**

**4 Loading the covariates** **4**

**5 Specifying the models in JAGS** **5**
- 5.1 Models without covariates . . . . . 5
  - 5.1.1 Ricker . . . . . 5
  - 5.1.2 Beverton-Holt . . . . . 8
- 5.2 Models with all covariates . . . . . 12
  - 5.2.1 Ricker . . . . . 12
  - 5.2.2 Beverton-Holt . . . . . 15

**6 Fitting the models** **18**
- 6.1 Models without covariates . . . . . 19
  - 6.1.1 Ricker . . . . . 20
  - 6.1.2 Beverton-Holt . . . . . 21
- 6.2 Models with all covariates . . . . . 22
  - 6.2.1 Ricker . . . . . 23
  - 6.2.2 Beverton-Holt . . . . . 23

**7 Model selection** **23**

**8 Model diagnostics** **25**


This is version 0.18.11.29.

#### 1 Background

This appendix describes how we fit the models and evaluated their relative performances. It demonstrates how to load the fish data and environmenal covariates, specify the different models in the **JAGS** software, and fit each one.

All analyses require the R software (v3.4.3 or later) for data retrieval, data processing, and summarizing model results, and the JAGS software (v4.2.0) for Markov chain Monte Carlo (MCMC) simulation. Please note that some of the **R** code below may not work with older versions of **JAGS** due to some changes in the ways that arrays are handled.

We also need a few packages that are not included with the base installation of **R**, so we begin by installing them (if necessary) and then loading them.

```
if(!require("here")) {
  install.packages("here")
  library("here")
}
if(!require("readr")) {
  install.packages("readr")
  library("readr")
}
if(!require("rjags")) {
  install.packages("rjags")
  library("rjags")
}
if(!require("loo")) {
  install.packages("loo")
  library("loo")
}
if(!require("ggplot2")) {
  install.packages("ggplot2")
  library("ggplot2")
}
## set directory locations
datadir <- here("data")
jagsdir <- here("jags")
analdir <- here("analysis")
savedir <- here("analysis/cache")
```

We also need a couple of helper functions.

```
## better round
Re2prec <- function(x, fun = "round", prec = 1) {
  ## 'fun' can be "round", "floor", or "ceiling"
  ## 'prec' is nearest value
  ## (eg, 0.1 is to nearest tenth; 1 is to nearest integer)
  if(prec<=0) { stop("\"prec\" cannot be less than or equal to 0") }
  do.call(map,list(x/prec))*prec
}

## wrapper function to fit JAGS models & rearrange output
fit_jags <- function(model, data, params, inits, ctrl, dir = jagsdir) {
  jm <- jags.model(file.path(jagsdir, model),
                   data,
                   inits,
                   ctrl$chains,
                   ctrl$burn,
```

```
                   quiet = TRUE)
  return(coda.samples(jm, params, ctrl$length, ctrl$thin))
}
```

#### 2 User inputs

We begin by supplying values for the following parameters, which we need for model fitting and evaluation.

```
## first & last years of fish data
yr_frst <- 1978
yr_last <- 2018

## min & max adult age classes
age_min <- 3
age_max <- 8
## years (if any) of age-comp to skip; see below
age_skip <- 0

## number of years for run forecasts
n_fore <- 0

## upper threshold for Gelman & Rubin's potential scale reduction factor (Rhat).
Rhat_thresh <- 1.1
```

Next we specify the names of three necessary data files containing the following information:

1. observed total number of adult spawners (escapement) by year;
2. observed age composition of adult spawners by year;
3. observed total harvest by year;

```
## 1. file with escapement data
## [n_yrs x 2] matrix of obs counts; 1st col is calendar yr
fn_esc <- "skagit_sthd_esc.csv"

## 2. file with age comp data
## [n_yrs x (1+A)]; 1st col is calendar yr
fn_age <- "skagit_sthd_age.csv"

## 3. file with harvest data
## [n_yrs x 2] matrix of obs catch; 1st col is calendar yr
fn_harv <- "skagit_sthd_catch.csv"
```

#### 3 Loading the fish data

Here we load in the first three data files and do some simple calculations and manipulations. First the spawner data:

```
## escapement
dat_esc <- read_csv(file.path(datadir, fn_esc))
## years of data
dat_yrs <- dat_esc$year
## number of years of data
n_yrs <- length(dat_yrs)
## log of escapement
ln_dat_esc <- c(log(dat_esc$escapement),rep(NA,n_fore))
```

Next the age composition data:

```
## age comp data
dat_age <- read_csv(file.path(datadir, fn_age))
## drop year col & first (age_min + age_skip) rows
dat_age <- dat_age[-(1:(age_min+age_skip)),-1]
## num of age classes
A <- age_max - age_min + 1
## add row(s) of NA's for forecast years
if(n_fore > 0) {
  dat_age <- rbind(dat_age,
                   matrix(0, n_fore, A,
                          dimnames = list(n_yrs+seq(n_fore),
                                          colnames(dat_age))))
}
## total num of age obs by cal yr
dat_age[,"sum"] <- apply(dat_age, 1, sum)
## row indices for any years with no obs age comp
idx_NA_yrs <- which(dat_age$sum<A, TRUE)
## replace 0's in yrs w/o any obs with NA's
dat_age[idx_NA_yrs,(1:A)] <- NA
## change total in yrs w/o any obs from 0 to A to help dmulti()
dat_age[idx_NA_yrs,"sum"] <- A
## convert class
dat_age <- as.matrix(dat_age)
```

And then the harvest data:

```
## harvest
dat_harv <- read_csv(file.path(datadir, fn_harv))
## drop year col & first age_max rows
dat_harv <- c(dat_harv$catch, rep(0,n_fore))
```

### 4 Loading the covariates

Our analysis investigates the effects of 4 covariates on the population's instrinic growth rate:

1. Maximum river discharge in winter;
2. Minimum river discharge in summer;
3. North Pacific Gyre Oscillation;

4. Releases of hatchery-born juveniles.

All of the covariates are contained in the file `/data/skagit_sthd_covars.csv`. We will load and then standardize them to have zero-mean and unit-variance.

```
## covariate(s)
dat_cvrs <- read_csv(file.path(datadir, "skagit_sthd_covars.csv"))
## drop year col
dat_cvrs <- dat_cvrs[,-1]
## transform the covariates to z-scores
scl_cvrs <- as.matrix(scale(dat_cvrs))
## total number of covariates
n_cov <- dim(scl_cvrs)[2]
```

### 5 Specifying the models in JAGS

Now we can specify the various models in JAGS. We fit a total of 4 different models, which we outline below, based on the 2 different process models with and without and covariates.

#### 5.1 Models without covariates

##### 5.1.1 Ricker

```
cat("

model {

  ##--------
  ## PRIORS
  ##--------
  ## alpha = exp(a) = intrinsic productivity
  alpha ~ dnorm(0,0.01) T(0,);
  mu_Rkr_a <- log(alpha);
  E_Rkr_a <- mu_Rkr_a + sigma_r/(2 - 2*phi^2);

  ## strength of dens depend
  beta_inv ~ dnorm(0, 1e-9) T(0,);
  beta <- 1/beta_inv;

  ## AR(1) coef for proc errors
  phi ~ dunif(-0.999,0.999);

  ## process variance for recruits model
  sigma_r ~ dnorm(0, 2e-2) T(0,);
  tau_r <- 1/sigma_r;

  ## innovation in first year
```

```
innov_1 ~ dnorm(0,tau_r*(1-phi*phi));

## obs variance for spawners
tau_s <- 1/sigma_s;
sigma_s ~ dnorm(0, 0.001) T(0,);

## maturity schedule
## unif vec for Dirch prior
theta <- c(1,10,10,5,1,1)
## hyper-mean for maturity
pi_eta ~ ddirch(theta);
## hyper-prec for maturity
pi_tau ~ dnorm(0, 0.01) T(0,);
for(t in 1:(n_yrs-age_min+n_fore)) { pi_vec[t,1:A] ~ ddirch(pi_eta*pi_tau) }

## unprojectable early recruits;
## hyper mean across all popns
Rec_mu ~ dnorm(0,0.001);
## hyper SD across all popns
Rec_sig ~ dunif(0,100);
## precision across all popns
Rec_tau <- pow(Rec_sig,-2);
## multipliers for unobservable total runs
  ttl_run_mu ~ dunif(1,5);
  ttl_run_tau ~ dunif(1,20);

## get total cal yr returns for first age_min yrs
for(i in 1:(age_min+age_skip)) {
      ln_tot_Run[i] ~ dnorm(ttl_run_mu*Rec_mu,Rec_tau/ttl_run_tau);
      tot_Run[i] <- exp(ln_tot_Run[i]);
}

## estimated harvest rate
for(t in 1:(n_yrs+n_fore)) { h_rate[t] ~ dunif(0,1) }

##------------
## LIKELIHOOD
##------------
## 1st brood yr requires different innovation
## predicted recruits in BY t
ln_Rkr_a[1] <- mu_Rkr_a;
E_ln_Rec[1] <- ln_Rkr_a[1] + ln_Sp[1] - beta*Sp[1] + phi*innov_1;
tot_ln_Rec[1] ~ dnorm(E_ln_Rec[1],tau_r);
res_ln_Rec[1] <- tot_ln_Rec[1] - E_ln_Rec[1];
## median of total recruits
tot_Rec[1] <- exp(tot_ln_Rec[1]);

## R/S
```

```
ln_RS[1] <- tot_ln_Rec[1] - ln_Sp[1];

## brood-yr recruits by age
for(a in 1:A) {
  Rec[1,a] <- tot_Rec[1] * pi_vec[1,a];
}

## brood years 2:(n_yrs-age_min)
for(t in 2:(n_yrs-age_min+n_fore)) {
  ## predicted recruits in BY t
  ln_Rkr_a[t] <- mu_Rkr_a;
  E_ln_Rec[t] <- ln_Rkr_a[t] + ln_Sp[t] - beta*Sp[t] + phi*res_ln_Rec[t-1];
  tot_ln_Rec[t] ~ dnorm(E_ln_Rec[t],tau_r);
  res_ln_Rec[t] <- tot_ln_Rec[t] - E_ln_Rec[t];
  ## median of total recruits
  tot_Rec[t] <- exp(tot_ln_Rec[t]);
  ## R/S
  ln_RS[t] <- tot_ln_Rec[t] - ln_Sp[t];
  ## brood-yr recruits by age
  for(a in 1:A) {
    Rec[t,a] <- tot_Rec[t] * pi_vec[t,a];
  }
} ## end t loop over year

## get predicted calendar year returns by age
## matrix Run has dim [(n_yrs-age_min) x A]
## step 1: incomplete early broods
## first cal yr of this grp is first brood yr + age_min + age_skip
for(i in 1:(age_max-age_min-age_skip)) {
  ## projected recruits
  for(a in 1:(i+age_skip)) {
    Run[i,a] <- Rec[(age_skip+i)-a+1,a];
  }
  ## imputed recruits
  for(a in (i+1+age_skip):A) {
    lnRec[i,a] ~ dnorm(Rec_mu,Rec_tau);
    Run[i,a] <- exp(lnRec[i,a]);
  }
  ## total run size
  tot_Run[i+age_min+age_skip] <- sum(Run[i,1:A]);
  ## predicted age-prop vec for multinom
  for(a in 1:A) {
    age_v[i,a] <- Run[i,a] / tot_Run[i+age_min];
  }
  ## multinomial for age comp
  dat_age[i,1:A] ~ dmulti(age_v[i,1:A],dat_age[i,A+1]);
  lp_age[i] <- logdensity.multi(dat_age[i,1:A],age_v[i,1:A],dat_age[i,A+1]);
}
```

```
  ## step 2: info from complete broods
  ## first cal yr of this grp is first brood yr + age_max
  for(i in (A-age_skip):(n_yrs-age_min-age_skip+n_fore)) {
    for(a in 1:A) {
      Run[i,a] <- Rec[(age_skip+i)-a+1,a];
    }
    ## total run size
    tot_Run[i+age_min+age_skip] <- sum(Run[i,1:A]);
    ## predicted age-prop vec for multinom
    for(a in 1:A) {
      age_v[i,a] <- Run[i,a] / tot_Run[i+age_min];
    }
    ## multinomial for age comp
    dat_age[i,1:A] ~ dmulti(age_v[i,1:A],dat_age[i,A+1]);
    lp_age[i] <- logdensity.multi(dat_age[i,1:A],age_v[i,1:A],dat_age[i,A+1]);
  }

  ## get predicted calendar year spawners
  ## first cal yr is first brood yr
  for(t in 1:(n_yrs+n_fore)) {
    ## obs model for spawners
    # Sp[t] <- max(10,tot_Run[t] - dat_harv[t]);
    est_harv[t] = h_rate[t] * tot_Run[t];
    dat_harv[t] ~ dlnorm(log(est_harv[t]), 20);
    Sp[t] = tot_Run[t] - est_harv[t];
    ln_Sp[t] <- log(Sp[t]);
    ln_dat_esc[t] ~ dnorm(ln_Sp[t], tau_s);
    lp_esc[t] <- logdensity.norm(ln_dat_esc[t],ln_Sp[t], tau_s);
  }

} ## end model description

", file=file.path(jagsdir, "IPM_RK_AR.txt"))
```

##### 5.1.2 Beverton-Holt

```
cat("

model {

  ##--------
  ## PRIORS
  ##--------
  ## alpha = exp(a) = intrinsic productivity
  alpha ~ dnorm(0,0.001) T(0,);
  mu_BH_a <- log(alpha);
```

```
E_BH_a <- mu_BH_a + sigma_r/(2 - 2*phi^2);

## strength of dens depend
beta_inv ~ dnorm(0, 1e-9) T(0,);
beta <- 1/beta_inv;

## AR(1) coef for proc errors
phi ~ dunif(-0.999,0.999);

## process variance for recruits model
sigma_r ~ dnorm(0, 2e-2) T(0,);
tau_r <- 1/sigma_r;

## innovation in first year
innov_1 ~ dnorm(0,tau_r*(1-phi*phi));

## obs variance for spawners
tau_s <- 1/sigma_s;
sigma_s ~ dnorm(0, 0.001) T(0,);

## unprojectable early recruits;
## hyper mean across all popns
Rec_mu ~ dnorm(0,0.001);
## hyper SD across all popns
Rec_sig ~ dunif(0,100);
## precision across all popns
Rec_tau <- pow(Rec_sig,-2);
## multipliers for unobservable total runs
  ttl_run_mu ~ dunif(1,5);
  ttl_run_tau ~ dunif(1,20);

## get total cal yr returns for first age_min yrs
for(i in 1:(age_min+age_skip)) {
      ln_tot_Run[i] ~ dnorm(ttl_run_mu*Rec_mu,Rec_tau/ttl_run_tau);
      tot_Run[i] <- exp(ln_tot_Run[i]);
}

## maturity schedule
## unif vec for Dirch prior
theta <- c(1,10,10,5,1,1)
## hyper-mean for maturity
pi_eta ~ ddirch(theta);
## hyper-prec for maturity
pi_tau ~ dnorm(0, 0.01) T(0,);
for(t in 1:(n_yrs-age_min+n_fore)) { pi_vec[t,1:A] ~ ddirch(pi_eta*pi_tau) }

## estimated harvest rate
for(t in 1:(n_yrs+n_fore)) { h_rate[t] ~ dunif(0,1) }
```

```
##------------
## LIKELIHOOD
##------------
## 1st brood yr requires different innovation
## predicted recruits in BY t
ln_BH_a[1] <- mu_BH_a;
E_ln_Rec[1] <- ln_BH_a[1] + ln_Sp[1] - log(1 + beta*Sp[1]) + phi*innov_1;
tot_ln_Rec[1] ~ dnorm(E_ln_Rec[1],tau_r);
res_ln_Rec[1] <- tot_ln_Rec[1] - E_ln_Rec[1];
## median of total recruits
tot_Rec[1] <- exp(tot_ln_Rec[1]);

## R/S
ln_RS[1] <- tot_ln_Rec[1] - ln_Sp[1];

## brood-yr recruits by age
for(a in 1:A) {
  Rec[1,a] <- tot_Rec[1] * pi_vec[1,a];
}

## brood years 2:(n_yrs-age_min)
for(t in 2:(n_yrs-age_min+n_fore)) {
  ## predicted recruits in BY t
  ln_BH_a[t] <- mu_BH_a;
  E_ln_Rec[t] <- ln_BH_a[t] + ln_Sp[t] - log(1 + beta*Sp[t]) + phi*res_ln_Rec[t-1];
  tot_ln_Rec[t] ~ dnorm(E_ln_Rec[t],tau_r);
  res_ln_Rec[t] <- tot_ln_Rec[t] - E_ln_Rec[t];
  ## median of total recruits
  tot_Rec[t] <- exp(tot_ln_Rec[t]);
  ## R/S
  ln_RS[t] <- tot_ln_Rec[t] - ln_Sp[t];
  ## brood-yr recruits by age
  for(a in 1:A) {
    Rec[t,a] <- tot_Rec[t] * pi_vec[t,a];
  }
} ## end t loop over year

## get predicted calendar year returns by age
## matrix Run has dim [(n_yrs-age_min) x A]
## step 1: incomplete early broods
## first cal yr of this grp is first brood yr + age_min + age_skip
for(i in 1:(age_max-age_min-age_skip)) {
  ## projected recruits
  for(a in 1:(i+age_skip)) {
    Run[i,a] <- Rec[(age_skip+i)-a+1,a];
  }
  ## imputed recruits
```

```
    for(a in (i+1+age_skip):A) {
      lnRec[i,a] ~ dnorm(Rec_mu,Rec_tau);
      Run[i,a] <- exp(lnRec[i,a]);
    }
    ## total run size
    tot_Run[i+age_min+age_skip] <- sum(Run[i,1:A]);
    ## predicted age-prop vec for multinom
    for(a in 1:A) {
      age_v[i,a] <- Run[i,a] / tot_Run[i+age_min];
    }
    ## multinomial for age comp
    dat_age[i,1:A] ~ dmulti(age_v[i,1:A],dat_age[i,A+1]);
    lp_age[i] <- logdensity.multi(dat_age[i,1:A],age_v[i,1:A],dat_age[i,A+1]);
  }

  ## step 2: info from complete broods
  ## first cal yr of this grp is first brood yr + age_max
  for(i in (A-age_skip):(n_yrs-age_min-age_skip+n_fore)) {
    for(a in 1:A) {
      Run[i,a] <- Rec[(age_skip+i)-a+1,a];
    }
    ## total run size
    tot_Run[i+age_min+age_skip] <- sum(Run[i,1:A]);
    ## predicted age-prop vec for multinom
    for(a in 1:A) {
      age_v[i,a] <- Run[i,a] / tot_Run[i+age_min];
    }
    ## multinomial for age comp
    dat_age[i,1:A] ~ dmulti(age_v[i,1:A],dat_age[i,A+1]);
    lp_age[i] <- logdensity.multi(dat_age[i,1:A],age_v[i,1:A],dat_age[i,A+1]);
  }

  ## get predicted calendar year spawners
  ## first cal yr is first brood yr
  for(t in 1:(n_yrs+n_fore)) {
    ## obs model for spawners
    # Sp[t] <- max(10,tot_Run[t] - dat_harv[t]);
    est_harv[t] = h_rate[t] * tot_Run[t];
    dat_harv[t] ~ dlnorm(log(est_harv[t]), 20);
    Sp[t] = tot_Run[t] - est_harv[t];
    ln_Sp[t] <- log(Sp[t]);
    ln_dat_esc[t] ~ dnorm(ln_Sp[t], tau_s);
    lp_esc[t] <- logdensity.norm(ln_dat_esc[t],ln_Sp[t], tau_s);
  }

} ## end model description

", file=file.path(jagsdir, "IPM_BH_AR.txt"))
```

##### 5.2 Models with all covariates

###### 5.2.1 Ricker

```
cat("

model {

  ##--------
  ## PRIORS
  ##--------
  ## alpha = exp(a) = intrinsic productivity
  alpha ~ dnorm(0,0.01) T(0,);
  mu_Rkr_a <- log(alpha);
  E_Rkr_a <- mu_Rkr_a + sigma_r/(2 - 2*phi^2);

  ## strength of dens depend
  beta_inv ~ dnorm(0, 1e-9) T(0,);
  beta <- 1/beta_inv;

  ## covariate effects
  for(i in 1:n_cov) { gamma[i] ~ dnorm(0,0.01) }

  ## AR(1) coef for proc errors
  # phi ~ dunif(-0.999,0.999);
  phi <- 0;

  ## process variance for recruits model
  sigma_r ~ dnorm(0, 2e-2) T(0,);
  tau_r <- 1/sigma_r;

  ## innovation in first year
  innov_1 ~ dnorm(0,tau_r*(1-phi*phi));

  ## obs variance for spawners
  tau_s <- 1/sigma_s;
  sigma_s ~ dnorm(0, 0.001) T(0,);

  ## maturity schedule
  ## unif vec for Dirch prior
  theta <- c(1,10,10,5,1,1)
  ## hyper-mean for maturity
  pi_eta ~ ddirch(theta);
  ## hyper-prec for maturity
  pi_tau ~ dnorm(0, 0.01) T(0,);
  for(t in 1:(n_yrs-age_min+n_fore)) { pi_vec[t,1:A] ~ ddirch(pi_eta*pi_tau) }
```

```
## unprojectable early recruits;
## hyper mean across all popns
Rec_mu ~ dnorm(0,0.001);
## hyper SD across all popns
Rec_sig ~ dunif(0,100);
## precision across all popns
Rec_tau <- pow(Rec_sig,-2);
## multipliers for unobservable total runs
  ttl_run_mu ~ dunif(1,5);
  ttl_run_tau ~ dunif(1,20);

## get total cal yr returns for first age_min yrs
for(i in 1:(age_min+age_skip)) {
      ln_tot_Run[i] ~ dnorm(ttl_run_mu*Rec_mu,Rec_tau/ttl_run_tau);
      tot_Run[i] <- exp(ln_tot_Run[i]);
}

## estimated harvest rate
for(t in 1:(n_yrs+n_fore)) { h_rate[t] ~ dunif(0,1) }

##------------
## LIKELIHOOD
##------------
## 1st brood yr requires different innovation
## predicted recruits in BY t
covar[1] <- inprod(gamma,mod_cvrs[1,]);
ln_Rkr_a[1] <- mu_Rkr_a + covar[1];
E_ln_Rec[1] <- ln_Rkr_a[1] + ln_Sp[1] - beta*Sp[1] + phi*innov_1;
tot_ln_Rec[1] ~ dnorm(E_ln_Rec[1],tau_r);
res_ln_Rec[1] <- tot_ln_Rec[1] - E_ln_Rec[1];
## median of total recruits
tot_Rec[1] <- exp(tot_ln_Rec[1]);

## R/S
ln_RS[1] <- tot_ln_Rec[1] - ln_Sp[1];

## brood-yr recruits by age
for(a in 1:A) {
  Rec[1,a] <- tot_Rec[1] * pi_vec[1,a];
}

## brood years 2:(n_yrs-age_min)
for(t in 2:(n_yrs-age_min+n_fore)) {
  ## predicted recruits in BY t
  covar[t] <- inprod(gamma, mod_cvrs[t,]);
  ln_Rkr_a[t] <- mu_Rkr_a + covar[t];
  E_ln_Rec[t] <- ln_Rkr_a[t] + ln_Sp[t] - beta*Sp[t] + phi*res_ln_Rec[t-1];
  tot_ln_Rec[t] ~ dnorm(E_ln_Rec[t],tau_r);
```

```
  res_ln_Rec[t] <- tot_ln_Rec[t] - E_ln_Rec[t];
  ## median of total recruits
  tot_Rec[t] <- exp(tot_ln_Rec[t]);
  ## R/S
  ln_RS[t] <- tot_ln_Rec[t] - ln_Sp[t];
  ## brood-yr recruits by age
  for(a in 1:A) {
    Rec[t,a] <- tot_Rec[t] * pi_vec[t,a];
  }
} ## end t loop over year

## get predicted calendar year returns by age
## matrix Run has dim [(n_yrs-age_min) x A]
## step 1: incomplete early broods
## first cal yr of this grp is first brood yr + age_min + age_skip
for(i in 1:(age_max-age_min-age_skip)) {
  ## projected recruits
  for(a in 1:(i+age_skip)) {
    Run[i,a] <- Rec[(age_skip+i)-a+1,a];
  }
  ## imputed recruits
  for(a in (i+1+age_skip):A) {
    lnRec[i,a] ~ dnorm(Rec_mu,Rec_tau);
    Run[i,a] <- exp(lnRec[i,a]);
  }
  ## total run size
  tot_Run[i+age_min+age_skip] <- sum(Run[i,1:A]);
  ## predicted age-prop vec for multinom
  for(a in 1:A) {
    age_v[i,a] <- Run[i,a] / tot_Run[i+age_min];
  }
  ## multinomial for age comp
  dat_age[i,1:A] ~ dmulti(age_v[i,1:A],dat_age[i,A+1]);
  lp_age[i] <- logdensity.multi(dat_age[i,1:A],age_v[i,1:A],dat_age[i,A+1]);
}

## step 2: info from complete broods
## first cal yr of this grp is first brood yr + age_max
for(i in (A-age_skip):(n_yrs-age_min-age_skip+n_fore)) {
  for(a in 1:A) {
    Run[i,a] <- Rec[(age_skip+i)-a+1,a];
  }
  ## total run size
  tot_Run[i+age_min+age_skip] <- sum(Run[i,1:A]);
  ## predicted age-prop vec for multinom
  for(a in 1:A) {
    age_v[i,a] <- Run[i,a] / tot_Run[i+age_min];
  }
```

```
    ## multinomial for age comp
    dat_age[i,1:A] ~ dmulti(age_v[i,1:A],dat_age[i,A+1]);
    lp_age[i] <- logdensity.multi(dat_age[i,1:A],age_v[i,1:A],dat_age[i,A+1]);
  }

  ## get predicted calendar year spawners
  ## first cal yr is first brood yr
  for(t in 1:(n_yrs+n_fore)) {
    ## obs model for spawners
    # Sp[t] <- max(10,tot_Run[t] - dat_harv[t]);
    est_harv[t] = h_rate[t] * tot_Run[t];
    dat_harv[t] ~ dlnorm(log(est_harv[t]), 20);
    Sp[t] = tot_Run[t] - est_harv[t];
    ln_Sp[t] <- log(Sp[t]);
    ln_dat_esc[t] ~ dnorm(ln_Sp[t], tau_s);
    lp_esc[t] <- logdensity.norm(ln_dat_esc[t],ln_Sp[t], tau_s);
  }

} ## end model description

", file=file.path(jagsdir, "IPM_RK_cov_AR.txt"))
```

##### 5.2.2 Beverton-Holt

```
cat("

model {

  ##--------
  ## PRIORS
  ##--------
  ## alpha = intrinsic productivity
  alpha ~ dnorm(0,0.001) T(0,);
  mu_BH_a <- log(alpha);
  E_BH_a <- mu_BH_a + sigma_r/(2 - 2*phi^2);

  ## strength of dens depend
  beta_inv ~ dnorm(0, 1e-9) T(0,);
  beta <- 1/beta_inv;

  ## covariate effects
  for(i in 1:n_cov) { gamma[i] ~ dnorm(0,0.01) }

  ## AR(1) coef for proc errors
  # phi ~ dunif(-0.999,0.999);
  phi <- 0;
```

```
## innovation in first year
innov_1 ~ dnorm(0,tau_r*(1-phi*phi));

## process variance for recruits model
sigma_r ~ dnorm(0, 2e-2) T(0,);
tau_r <- 1/sigma_r;

## obs variance for spawners
tau_s <- 1/sigma_s;
sigma_s ~ dnorm(0, 0.001) T(0,);

## unprojectable early recruits;
## hyper mean across all popns
Rec_mu ~ dnorm(0,0.001);
## hyper SD across all popns
Rec_sig ~ dunif(0,100);
## precision across all popns
Rec_tau <- pow(Rec_sig,-2);
## multipliers for unobservable total runs
  ttl_run_mu ~ dunif(1,5);
  ttl_run_tau ~ dunif(1,20);

## get total cal yr returns for first age_min yrs
for(i in 1:(age_min+age_skip)) {
      ln_tot_Run[i] ~ dnorm(ttl_run_mu*Rec_mu,Rec_tau/ttl_run_tau);
      tot_Run[i] <- exp(ln_tot_Run[i]);
}

## maturity schedule
## unif vec for Dirch prior
theta <- c(1,10,10,5,1,1)
## hyper-mean for maturity
pi_eta ~ ddirch(theta);
## hyper-prec for maturity
pi_tau ~ dnorm(0, 0.01) T(0,);
for(t in 1:(n_yrs-age_min+n_fore)) { pi_vec[t,1:A] ~ ddirch(pi_eta*pi_tau) }

## estimated harvest rate
for(t in 1:(n_yrs+n_fore)) { h_rate[t] ~ dunif(0,1) }

##------------
## LIKELIHOOD
##------------
## predicted recruits in BY t
covar[1] <- inprod(gamma,mod_cvrs[1,]);
ln_BH_a[1] <- mu_BH_a + covar[1];
E_ln_Rec[1] <- ln_BH_a[1] + ln_Sp[1] - log(1 + beta*Sp[1]) + phi*innov_1;
tot_ln_Rec[1] ~ dnorm(E_ln_Rec[1],tau_r);
```

```
res_ln_Rec[1] <- tot_ln_Rec[1] - E_ln_Rec[1];
## median of total recruits
tot_Rec[1] <- exp(tot_ln_Rec[1]);

## R/S
ln_RS[1] <- tot_ln_Rec[1] - ln_Sp[1];

## brood-yr recruits by age
for(a in 1:A) {
  Rec[1,a] <- tot_Rec[1] * pi_vec[1,a];
}

## brood years 2:(n_yrs-age_min)
for(t in 2:(n_yrs-age_min+n_fore)) {
  ## predicted recruits in BY t
  covar[t] <- inprod(gamma, mod_cvrs[t,]);
  ln_BH_a[t] <- mu_BH_a + covar[t];
  E_ln_Rec[t] <- ln_BH_a[t] + ln_Sp[t] - log(1 + beta*Sp[t]) + phi*res_ln_Rec[t-1];
  tot_ln_Rec[t] ~ dnorm(E_ln_Rec[t],tau_r);
  res_ln_Rec[t] <- tot_ln_Rec[t] - E_ln_Rec[t];
  ## median of total recruits
  tot_Rec[t] <- exp(tot_ln_Rec[t]);
  ## R/S
  ln_RS[t] <- tot_ln_Rec[t] - ln_Sp[t];
  ## brood-yr recruits by age
  for(a in 1:A) {
    Rec[t,a] <- tot_Rec[t] * pi_vec[t,a];
  }
} ## end t loop over year

## get predicted calendar year returns by age
## matrix Run has dim [(n_yrs-age_min) x A]
## step 1: incomplete early broods
## first cal yr of this grp is first brood yr + age_min + age_skip
for(i in 1:(age_max-age_min-age_skip)) {
  ## projected recruits
  for(a in 1:(i+age_skip)) {
    Run[i,a] <- Rec[(age_skip+i)-a+1,a];
  }
  ## imputed recruits
  for(a in (i+1+age_skip):A) {
    lnRec[i,a] ~ dnorm(Rec_mu,Rec_tau);
    Run[i,a] <- exp(lnRec[i,a]);
  }
  ## total run size
  tot_Run[i+age_min+age_skip] <- sum(Run[i,1:A]);
  ## predicted age-prop vec for multinom
  for(a in 1:A) {
```

```
      age_v[i,a] <- Run[i,a] / tot_Run[i+age_min];
    }
    ## multinomial for age comp
    dat_age[i,1:A] ~ dmulti(age_v[i,1:A],dat_age[i,A+1]);
    lp_age[i] <- logdensity.multi(dat_age[i,1:A],age_v[i,1:A],dat_age[i,A+1]);
  }

  ## step 2: info from complete broods
  ## first cal yr of this grp is first brood yr + age_max
  for(i in (A-age_skip):(n_yrs-age_min-age_skip+n_fore)) {
    for(a in 1:A) {
      Run[i,a] <- Rec[(age_skip+i)-a+1,a];
    }
    ## total run size
    tot_Run[i+age_min+age_skip] <- sum(Run[i,1:A]);
    ## predicted age-prop vec for multinom
    for(a in 1:A) {
      age_v[i,a] <- Run[i,a] / tot_Run[i+age_min];
    }
    ## multinomial for age comp
    dat_age[i,1:A] ~ dmulti(age_v[i,1:A],dat_age[i,A+1]);
    lp_age[i] <- logdensity.multi(dat_age[i,1:A],age_v[i,1:A],dat_age[i,A+1]);
  }

  ## get predicted calendar year spawners
  ## first cal yr is first brood yr
  for(t in 1:(n_yrs+n_fore)) {
    ## obs model for spawners
    # Sp[t] <- max(10,tot_Run[t] - dat_harv[t]);
    est_harv[t] = h_rate[t] * tot_Run[t];
    dat_harv[t] ~ dlnorm(log(est_harv[t]), 20);
    Sp[t] = tot_Run[t] - est_harv[t];
    ln_Sp[t] <- log(Sp[t]);
    ln_dat_esc[t] ~ dnorm(ln_Sp[t], tau_s);
    lp_esc[t] <- logdensity.norm(ln_dat_esc[t],ln_Sp[t], tau_s);
  }

} ## end model description

", file=file.path(jagsdir, "IPM_BH_cov_AR.txt"))
```

### 6 Fitting the models

Before fitting the model in JAGS, we need to specify:

1. the data and indices that go into the model;
2. the model parameters and states that we want JAGS to return;

3. the MCMC control parameters.

```
## 1. Data to pass to JAGS:
dat_jags <- list(dat_age = dat_age,
                 ln_dat_esc = ln_dat_esc,
                 dat_harv = dat_harv,
                 A = A,
                 age_min = age_min,
                 age_max = age_max,
                 age_skip = age_skip,
                 n_yrs = n_yrs,
                 n_fore = n_fore)

## 2. Model params/states for JAGS to return:
##
##    These are specific to the process model,
##    so we define them in 'par_jags' below.

## 3. MCMC control params:
mcmc_ctrl <- list(
  chains = 4,
  length = 5e5,
  burn = 2e5,
  thin = 400
)
## total number of MCMC samples after burnin
mcmc_samp <- mcmc_ctrl$length*mcmc_ctrl$chains/mcmc_ctrl$thin
```

##### 6.1 Models without covariates

Please note that the following code takes ~80 min to run on a quad-core machine with 3.5 GHz Intel processors.

```
## empty list for fits
n_mods <- 4
mod_fits <- vector("list", n_mods)

## function for inits
init_vals_AR <- function() {
    list(alpha = 5,
         beta_inv = exp(mean(ln_dat_esc, na.rm = TRUE)),
         pi_tau = 10,
         pi_eta = rep(1,A),
         pi_vec = matrix(c(0.01,0.35,0.47,0.15,0.01,0.01),
                         n_yrs-age_min+n_fore, A,
                         byrow = TRUE),
         Rec_mu = log(1000),
         Rec_sig = 0.1,
```

```r
         tot_ln_Rec = rep(log(1000), n_yrs - age_min + n_fore),
         innov_1 = 0,
         phi = 0.5)
}
```

##### 6.1.1 Ricker

```r
## params/states to return
par_jags <- c("alpha","E_Rkr_a","mu_Rkr_a",
              "beta",
              "Sp","Rec","tot_ln_Rec","ln_RS",
              "pi_eta","pi_tau",
              "sigma_r","sigma_s","res_ln_Rec",
              "lp_age","lp_esc")
## fit model & save it
mod_fits[[1]] <- fit_jags("IPM_RK_AR.txt", dat_jags, par_jags, init_vals_AR, mcmc_ctrl)
```

###### 6.1.1.1 Convergence checks

```r
par_conv <- c("alpha", "beta",
              "sigma_r", "sigma_s",
              "pi_tau", paste0("pi_eta[",seq(A-1),"]"))

## Gelman-Rubin
gelman.diag(mod_fits[[1]][,par_conv])

## Potential scale reduction factors:
##
##            Point est. Upper C.I.
## alpha            1.04       1.11
## beta             1.04       1.13
## sigma_r          1.05       1.16
## sigma_s          1.01       1.02
## pi_tau           1.00       1.00
## pi_eta[1]        1.00       1.00
## pi_eta[2]        1.00       1.00
## pi_eta[3]        1.00       1.00
## pi_eta[4]        1.00       1.00
## pi_eta[5]        1.00       1.00
##
## Multivariate psrf
##
## 1.09

## autocorrelation
t(round(autocorr.diag(mod_fits[[1]][,par_conv],
                      lags = seq(mcmc_ctrl$thin, 4*mcmc_ctrl$thin, mcmc_ctrl$thin),
                      relative=FALSE), 2))
```

```
##           Lag 400 Lag 800 Lag 1200 Lag 1600
## alpha        0.10    0.05     0.05     0.04
## beta         0.13    0.08     0.07     0.05
## sigma_r      0.02    0.02     0.03     0.02
## sigma_s      0.24    0.20     0.17     0.19
## pi_tau       0.10    0.09     0.06     0.03
## pi_eta[1]    0.21    0.14     0.11     0.08
## pi_eta[2]    0.00    0.02     0.00     0.00
## pi_eta[3]   -0.02    0.00     0.03    -0.01
## pi_eta[4]    0.00    0.01    -0.01     0.01
## pi_eta[5]    0.15    0.08     0.06     0.03
```

##### 6.1.2 Beverton-Holt

```
## params/states to return
par_jags <- c("alpha","E_BH_a","mu_BH_a",
              "beta",
              "Sp","Rec","tot_ln_Rec","ln_RS",
              "pi_eta","pi_tau",
              "sigma_r","sigma_s","res_ln_Rec",
              "lp_age","lp_esc")
## fit model & save it
mod_fits[[2]] <- fit_jags("IPM_BH_AR.txt", dat_jags, par_jags, init_vals_AR, mcmc_ctrl)
```

###### 6.1.2.1 Convergence checks

```
## Gelman-Rubin
gelman.diag(mod_fits[[2]][,par_conv])
```

```
## Potential scale reduction factors:
##
##           Point est. Upper C.I.
## alpha           1.00       1.01
## beta            1.00       1.01
## sigma_r         1.00       1.00
## sigma_s         1.00       1.01
## pi_tau          1.02       1.05
## pi_eta[1]       1.00       1.00
## pi_eta[2]       1.00       1.00
## pi_eta[3]       1.00       1.00
## pi_eta[4]       1.00       1.00
## pi_eta[5]       1.00       1.01
##
## Multivariate psrf
##
## 1.18
```

```
## autocorrelation
t(round(autocorr.diag(mod_fits[[2]][,par_conv],
                      lags = seq(mcmc_ctrl$thin, 4*mcmc_ctrl$thin, mcmc_ctrl$thin),
                      relative=FALSE), 2))
```

```
##            Lag 400 Lag 800 Lag 1200 Lag 1600
## alpha         0.18    0.04     0.03     0.01
## beta          0.18    0.03     0.03     0.01
## sigma_r       0.02    0.01    -0.01    -0.02
## sigma_s       0.04    0.03    -0.03     0.03
## pi_tau        0.10    0.09     0.06     0.05
## pi_eta[1]     0.22    0.13     0.07     0.08
## pi_eta[2]     0.00    0.00     0.01     0.01
## pi_eta[3]     0.01   -0.01     0.00     0.00
## pi_eta[4]    -0.01    0.00    -0.02     0.04
## pi_eta[5]     0.15    0.09     0.02     0.01
```

##### 6.2 Models with all covariates

Now we fit the models that include covariates. Here is the `inits` function for `fit_jags()` and a specification of the covariates.

```
## function for inits
init_vals_cov <- function() {
  list(alpha = 5,
       beta_inv = exp(mean(ln_dat_esc, na.rm = TRUE)),
       gamma = rep(0, n_mods),
       pi_tau = 10,
       pi_eta = rep(1,A),
       pi_vec = matrix(c(0.01,0.35,0.47,0.15,0.01,0.01),
                       n_yrs-age_min+n_fore, A,
                       byrow = TRUE),
       Rec_mu = log(1000),
       Rec_sig = 0.1,
       tot_ln_Rec = rep(log(1000), n_yrs - age_min + n_fore),
       # phi = 0.5,
       innov_1 = 0)
}

## set of multi-covariate models
cset <- colnames(scl_cvrs)
dat_jags$n_cov <- length(cset)
dat_jags$mod_cvrs <- scl_cvrs[, cset]
```

##### 6.2.1 Ricker

```
## params/states to return
par_jags <- c("alpha","E_Rkr_a","mu_Rkr_a",
              "beta",
              "gamma",
              "Sp","Rec","tot_ln_Rec","ln_RS",
              "pi_eta","pi_tau",
              "sigma_r","sigma_s","res_ln_Rec",
              "lp_age","lp_esc")

## fit model & save it
mod_fits[[3]] <- fit_jags("IPM_RK_cov_AR.txt", dat_jags, par_jags,
                          init_vals_cov, mcmc_ctrl)
```

##### 6.2.2 Beverton-Holt

```
## params/states to return
par_jags <- c("alpha","E_BH_a","ln_BH_a",
              "beta",
              "gamma",
              "Sp","Rec","tot_ln_Rec","ln_RS",
              "pi_eta","pi_tau",
              "sigma_r","sigma_s","res_ln_Rec",
              "lp_age","lp_esc")

## fit model & save it
mod_fits[[4]] <- fit_jags("IPM_BH_cov_AR.txt", dat_jags, par_jags,
                          init_vals_cov, mcmc_ctrl)
```

### 7 Model selection

Via `loo()` and `compare()` with full table of results. Note that `elpd_diff` will be negative (positive) if the expected predictive accuracy for the first (second) model is higher.

```
LOOIC <- vector("list", n_mods)
## extract log densities from JAGS objects
for(i in 1:n_mods) {
  ## convert mcmc.list to matrix
  tmp_lp <- as.matrix(mod_fits[[i]])
  ## extract pointwise likelihoods
  tmp_lp <- tmp_lp[,grepl("lp_", colnames(tmp_lp))]
  ## if numerical underflows, convert -Inf to 5% less than min(likelihood)
  if(any(is.infinite(tmp_lp))) {
    tmp_lp[is.infinite(tmp_lp)] <- NA
    tmp_min <- min(tmp_lp, na.rm = TRUE)
```

```
    tmp_lp[is.na(tmp_lp)] <- tmp_min * 1.05
  }
  ## calculate LOOIC
  LOOIC[[i]] <- loo(tmp_lp)
}

## LOOIC for all data
tbl_LOOIC <- round(compare(x = LOOIC), 2)
rownames(tbl_LOOIC) <- sub("model", "", rownames(tbl_LOOIC))
tbl_LOOIC <- tbl_LOOIC[order(as.numeric(rownames(tbl_LOOIC))), ]
tbl_LOOIC <- cbind(model = rep(c("Ricker","B-H"),2),
                   covar = rep(c("No","Yes"),ea=2),
                   as.data.frame(tbl_LOOIC))
tbl_LOOIC[order(tbl_LOOIC[,"looic"]), ]
```

```
##    model covar elpd_diff elpd_loo se_elpd_loo  p_loo se_p_loo  looic se_looic
## 4    B-H   Yes      0.00  -379.21       48.16 136.66     9.63 758.42    96.32
## 2    B-H    No     -5.98  -385.19       50.09 153.59    14.34 770.39   100.18
## 3 Ricker   Yes    -19.13  -398.34       48.63 148.66    15.19 796.68    97.25
## 1 Ricker    No    -24.06  -403.27       50.83 159.91    19.85 806.53   101.66
```

```
## LOOIC for all data (pairwise)
## `elpd_diff` will be neg (pos) if the expected predictive accuracy
## for the first (second) model is higher.
## Ricker without vs with covariates
compare(LOOIC[[1]], LOOIC[[3]])
```

```
## elpd_diff        se
##       4.9      11.7
```

```
## BH without vs with covariates
compare(LOOIC[[2]], LOOIC[[4]])
```

```
## elpd_diff        se
##       6.0      10.3
```

```
## Ricker to BH (without covariates)
compare(LOOIC[[1]], LOOIC[[2]])
```

```
## elpd_diff        se
##      18.1       9.6
```

```
## Ricker to BH (with covariates)
compare(LOOIC[[3]], LOOIC[[4]])
```

```
## elpd_diff        se
##      19.1      10.8
```

```
## best model
best_i <- which(tbl_LOOIC[,"looic"] == min(tbl_LOOIC[,"looic"]))
best_fit <- mod_fits[[best_i]]
```

These results show that the Beverton-Holt model with covariates has the lowest LOOIC value. Therefore, the rest of the results will be based on that model.

### 8 Model diagnostics

Here is a table of the Gelman & Rubin statistics ($R_{hat}$) for the estimated parameters. Recall that we set an upper threshold of 1.1, so values larger than that deserve some additional inspection.

```
## params of interest
par_conv <- c("alpha","beta",paste0("gamma[",seq(4),"]"),
              "sigma_r","sigma_s","pi_tau",paste0("pi_eta[",seq(A-1),"]"))
## Gelman-Rubin
gelman.diag(best_fit[,par_conv])
```

```
## Potential scale reduction factors:
##
##              Point est. Upper C.I.
## alpha              1.01       1.02
## beta               1.01       1.02
## gamma[1]           1.00       1.00
## gamma[2]           1.00       1.00
## gamma[3]           1.00       1.00
## gamma[4]           1.00       1.00
## sigma_r            1.00       1.00
## sigma_s            1.01       1.01
## pi_tau             1.01       1.04
## pi_eta[1]          1.00       1.01
## pi_eta[2]          1.00       1.00
## pi_eta[3]          1.00       1.00
## pi_eta[4]          1.00       1.00
## pi_eta[5]          1.00       1.01
##
## Multivariate psrf
##
## 1.16
```

```
## Autocorrelation
t(round(autocorr.diag(best_fit[,par_conv],
                      lags = seq(mcmc_ctrl$thin, 4*mcmc_ctrl$thin, mcmc_ctrl$thin),
                      relative=FALSE), 2))
```

```
##              Lag 400 Lag 800 Lag 1200 Lag 1600
## alpha           0.40    0.18     0.09     0.07
## beta            0.40    0.18     0.08     0.07
## gamma[1]       -0.02    0.01    -0.02    -0.02
## gamma[2]       -0.01   -0.02     0.02    -0.02
## gamma[3]        0.03    0.03     0.02     0.00
## gamma[4]       -0.02   -0.01    -0.01     0.00
```

```
## sigma_r     -0.01    0.00     0.01    -0.02
## sigma_s      0.07    0.03     0.02     0.03
## pi_tau       0.13    0.08     0.08     0.07
## pi_eta[1]    0.25    0.15     0.11     0.07
## pi_eta[2]    0.01    0.00     0.02    -0.01
## pi_eta[3]    0.00   -0.01     0.01     0.00
## pi_eta[4]    0.01   -0.01     0.00     0.02
## pi_eta[5]    0.14    0.08     0.04     0.05
```
