## Appendix 3 - Summarize results for "An integrated population model for estimating the relative effects of natural and anthropogenic factors on a threatened population of Pacific trout"

### Appendix S3. Steps to recreate figures from main text.

Supporting information for Scheuerell et al.

#### Contents

**1 Background** **1**

**2 Load the information** **2**

**3 Model forms** **3**

**4 Main results** **5**

This is version 0.19.07.09.

#### 1 Background

This appendix shows how to recreate the figures in the main text based on the results from the best of the fitted models. All analyses require the R software (v3.5 or later), as well as a few packages that are not included with the base installation of R.

```
if(!require("readr")) {
  install.packages("readr")
  library("readr")
}
if(!require("captioner")) {
  devtools::install_github("adletaw/captioner")
  library("captioner")
}
if(!require("coda")) {
  install.packages("coda")
  library("coda")
}
if(!require("here")) {
  install.packages("here")
  library("here")
}
if(!require("gsl")) {
  install.packages("gsl")
  library("gsl")
```

```
}

## set default caption delimter
fig_cap <- captioner(suffix = ".")

## set directory locations
datadir <- here("data")
analdir <- here("analysis")
savedir <- here("analysis/cache")

## better round/floor/ceiling
around <- function(x, func = "round", prec = 1) {
  ## `func` can be "round", "floor", or "ceiling"
  ## `prec` is desired precision (eg, 0.1 is to nearest tenth)
  if(!is.double(x)) {
    stop("`x` must be a real number")
  }
  if(!(func %in% c("round", "floor", "ceiling"))) {
    stop("`func` must be \"round\", \"floor\", or \"ceiling\"")
  }
  if(prec <= 0) {
    stop("`prec` cannot be less than or equal to 0")
  }
  do.call(func, list(x / prec)) * prec
}
```

### 2 Load the information

Here we load in the estimated parameters and states from the selected model, as well as the covariates and harvest data and escapement data.

```
best_fit <- readRDS(file.path(savedir, "fit_bh_cov.rds"))
```

```
## covariate(s)
dat_cvrs <- read_csv(file.path(datadir, "skagit_sthd_covars.csv"))
## total number of covariates
n_cov <- dim(dat_cvrs)[2] - 1
```

```
## escapement
dat_esc <- read_csv(file.path(datadir, "skagit_sthd_esc.csv"))
## log of escapement
ln_dat_esc <- c(log(dat_esc$escapement), rep(NA, n_fore))
```

```
## harvest
dat_harv <- read_csv(file.path(datadir, "skagit_sthd_catch.csv"))
## drop year col & first age_max rows
dat_harv <- c(dat_harv$catch, rep(0, n_fore))
```

#### 3 Model forms

Here are the model parameters we used for the schematics of the deterministic forms for the Ricker and Beverton-Holt models.

```
## params
## Ricker
ra <- 3
rb <- 1.2e-4
## B-H
ba <- 3
bb <- 3/1.4e4

## ref pts
## Ricker
rmr <- ra/rb*exp(-1)
rsy <- (1 - lambert_W0(exp(1)/ra)) / rb
ruy <- 1 - lambert_W0(exp(1)/ra)
## B-H
bmr <- ba/bb
bsy <- (ba/bb)*sqrt(1/ba)-(1/bb)
bsy <- (sqrt(ba)-1)/bb
buy <- 1 - sqrt(1/ba)

## S-R curves
## spawners
ss <- seq(0,1.2e4,10)
## recuits (Ricker)
rr <- ra*ss/exp(rb*ss)
## recruits (B-H)
br <- ba*ss/(1 + bb*ss)
```

Here is the code to recreate the schematics in Figure 1.

```
layout(matrix(c(1,0,2),3,1),
       heights=lcm(c(3,0.3,3)*2.54),
       widths=lcm(3*2.54))

par(mai=c(0.4,0.4,0.2,0.2), omi=c(0,0,0,0.25))

## Ricker
plot(ss, rr, type="n", xlim=range(ss), ylim=range(ss), xaxs="i", yaxs="i",
     xlab="", ylab="", xaxt="n", yaxt="n", bty="L")
mtext(expression(italic(S[t])), 1, line=1, cex=1.1, at=max(ss))
mtext(expression(italic(R[t])), 2, line=0.5, cex=1.1, at=max(ss), las=1)
rttl <- "(a) Ricker"
text(400, max(ss), rttl, cex=1.1, adj=c(0,1), xpd=NA)
## 1:1
abline(a=0, b=1, col="gray")
```

```
#text(1.2e4, 1.2e4, "1:1", adj=c(1,0))
## R-S
lines(ss, rr, lwd=2)
rmod <- expression(frac(italic(alpha * S[t]),italic(e^{beta * S[t]})))
text(12300, ra*max(ss)/exp(rb*max(ss)), rmod, adj=c(0,0.5), xpd=NA)
## alpha
segments(0, 0, 1900, ra*1900, lty="dashed")
text(2000, ra*2000, expression(alpha), adj=c(0.5,0.5))
## MSY
segments(rsy,0,rsy,ra*rsy/exp(rb*rsy), lty="dashed")
text(rsy, 0, expression(frac(1-italic(W)~bgroup("(",frac(italic(e),alpha),")"),beta)), adj=c(0
segments(par()$usr[1],ra*rsy/exp(rb*rsy),rsy,ra*rsy/exp(rb*rsy), lty="dashed")
text(0, ra*rsy/exp(rb*rsy), expression(italic(R)[MSY]), pos=2, xpd=NA)
## K
segments(0, log(ra)/rb, log(ra)/rb, log(ra)/rb, lty="dashed")
segments(log(ra)/rb, 0, log(ra)/rb, log(ra)/rb, lty="dashed")
text(log(ra)/rb, 0, expression(frac(log(alpha),beta)), adj=c(0.5,1.2), xpd=NA)
text(0, log(ra)/rb, expression(italic(K)), pos=2, xpd=NA)

## B-H
plot(ss, br, type="n", xlim=range(ss), ylim=range(ss), xaxs="i", yaxs="i",
     xlab="", ylab="", xaxt="n", yaxt="n", bty="L")
mtext(expression(italic(S[t])), 1, line=1, cex=1.1, at=max(ss))
mtext(expression(italic(R[t])), 2, line=0.5, cex=1.1, at=max(ss), las=1)
bttl <- "(b) Beverton-Holt"
text(400, max(ss), bttl, cex=1.1, adj=c(0,1), xpd=NA)
## 1:1
abline(a=0, b=1, col="gray")
## R-S
lines(ss, br, lwd=2)
bmod <- expression(frac(italic(alpha * S[t]),1+italic(beta * S[t])))
text(max(ss)+300, ba*max(ss)/(1 + bb*max(ss)), bmod, adj=c(0,0.5), xpd=NA)
## alpha
segments(0, 0, 1500, ba*1500, lty="dashed")
text(1600, ba*1600, expression(alpha), adj=c(0.5,0.5))
## MSY
segments(bsy,0,bsy,ba*bsy/(1 + bb*bsy), lty="dashed")
text(bsy, 0, expression(frac(root(alpha)-1,beta)), adj=c(0.5,1.2), xpd=NA)
segments(par()$usr[1],ba*bsy/(1 + bb*bsy),bsy,ba*bsy/(1 + bb*bsy), lty="dashed")
text(0, ba*bsy/(1 + bb*bsy), expression(italic(R)[MSY]), pos=2, xpd=NA)
## K
segments(0, (ba-1)/bb, (ba-1)/bb, (ba-1)/bb, lty="dashed")
segments((ba-1)/bb, 0, (ba-1)/bb, (ba-1)/bb, lty="dashed")
text((ba-1)/bb, 0, expression(frac(alpha-1,beta)), adj=c(0.5,1.2), xpd=NA)
text(0, (ba-1)/bb, expression(italic(K)), pos=2, xpd=NA)
```

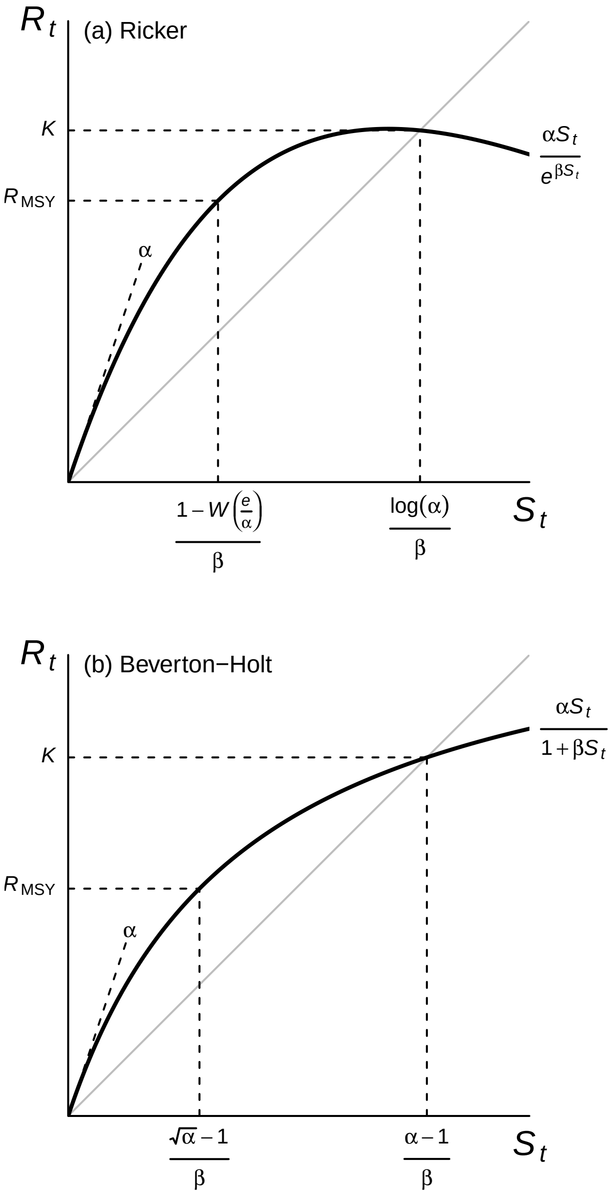

Figure 1. Deterministic forms of the (a) Ricker and (b) Beverton-Holt models used in the analyses (thick lines), including equations for carrying capacity ($K$) and the number of recruits corresponding to the maximum sustained yield ($R_{\text{MSY}}$). The parameter $\alpha$ defines the slope at the origin, the constant $e$ is Euler's number, and $W$ is the Lambert function (see Scheuerell 2016 for details). The gray line is where $R_t = S_t$.

### 4 Main results

We need to convert the `mcmc.list` output into a more user-friendly form for plotting, etc.

```
## results
mod_res <- do.call("rbind", best_fit)
```

#### 4.1 Total population size

Here is our estimate of the total run size (i.e., catch + escapement) over time. The black points are the data, the blue line is the median posterior estimate, and the shaded region is the 95% credible interval. Note that the y-axis is on a log scale.

```
clr <- rgb(0, 0, 255, alpha = 50, maxColorValue = 255)
## estimated spawner data for plotting
p_dat <- mod_res[,grep("Sp", colnames(mod_res))]
p_dat <- apply(p_dat, 2, quantile, CI_vec)
p_dat <- p_dat + matrix(dat_harv, length(CI_vec), n_yrs+n_fore, byrow = TRUE)
## time seq
t_idx_f <- seq(yr_frst, length.out = n_yrs+n_fore)
## plot
yp_min <- min(p_dat)
yp_max <- max(p_dat)
par(mai = c(0.8,0.8,0.1,0.1), omi = c(0.5,0.2,0.1,0.2))
plot(t_idx_f, p_dat[3,], ylim = c(yp_min,yp_max), type = "n",
     log = "y", xaxt = "n", yaxt = "n", bty = "L",
     xlab = "Year", ylab = "Run size (Catch + Escapement)", main = "", cex.lab = 1.2)
polygon(c(t_idx_f, rev(t_idx_f)), c(p_dat[3,], rev(p_dat[1,])),
        col = clr, border = NA)
lines(t_idx_f, p_dat[2,], col = "blue3", lwd = 2)
points(t_idx_f, exp(ln_dat_esc) + dat_harv, pch = 16, cex = 1)
axis(1, at = seq(1980, 2015, 5))
axis(2, at = c(4000, 8000, 16000))
```

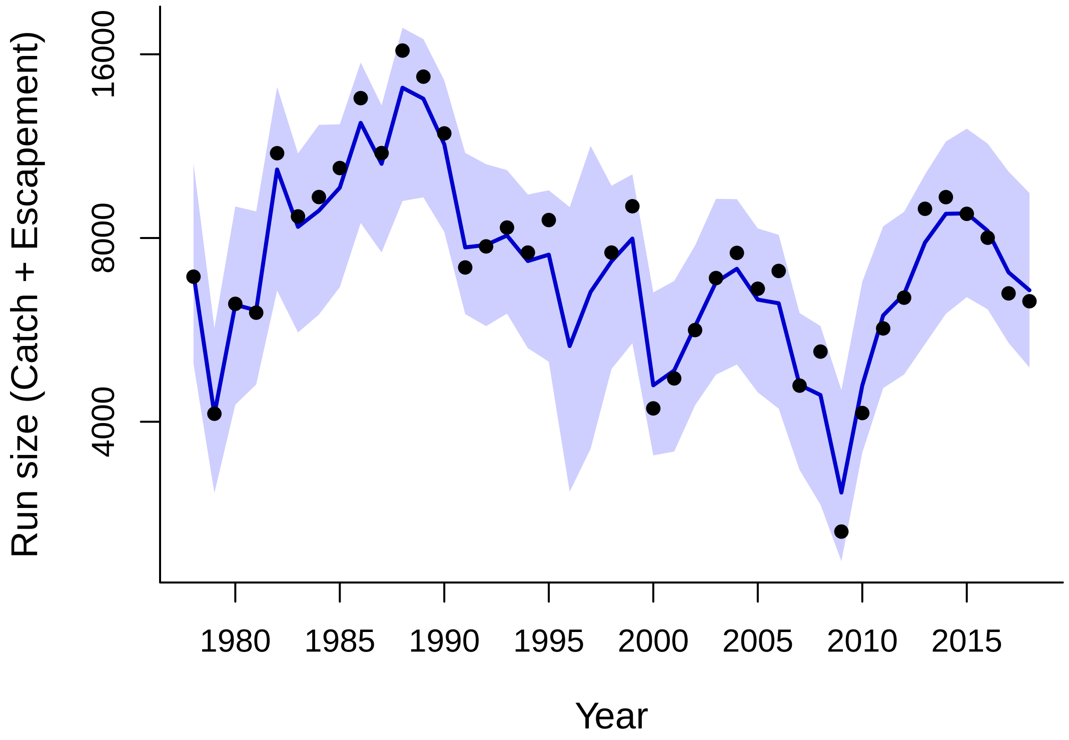

Figure 2. Time series of the estimated total population size (catch plus the adults that escaped to

spawn). The observed data are the points; the solid line is the median estimate and the shaded region indicates the 95% credible interval.

##### 4.2 Spawner-recruit relationship

Here is the relationship between spawner and subsequent recruits (a), assuming mean values for all covariates. Gray lines show the median relationship for each of the 41 years based on $a_t$. Note that for plotting purposes only in (b) and (c), the density in the largest bin for each parameter contains counts for all values greater or equal to that. Vertical arrows under the x-axes in (b) and (c) indicate the 2.5$^{\text{th}}$, 50$^{\text{th}}$, and 97.5$^{\text{th}}$ percentiles.

```
layout(matrix(c(1,1,2,3),2,2),c(3,2),c(1,1))
xoffSet <- 0.05
yoffSet <- 0.03

## colors for plotting
clr <- rgb(100, 0, 200,
           alpha = seq(200, 100,
                       length.out = age_max-age_min+n_fore),
           maxColorValue = 255)

## posterior of spawners
s_dat <- mod_res[,grep("Sp", colnames(mod_res))]
s_dat <- apply(s_dat, 2, quantile, CI_vec)
s_dat <- s_dat[, 1:(n_yrs-age_min+n_fore)]

## posterior of recruits
r_dat <- mod_res[, grep("tot_ln_Rec", colnames(mod_res))]
r_dat <- exp(apply(r_dat, 2, quantile, CI_vec))

## median values for a & b
aa <- apply(mod_res[, grep("ln_BH_a", colnames(mod_res))], 2, median)
bb <- median(mod_res[, "beta"])

## empty plot space for spawner-recruit relationships
dd <- 3000
yM <- around(max(r_dat), "ceiling", dd)
xM <- around(max(s_dat), "ceiling", dd)
par(mai = c(0.8,0.8,0.1,0.1), omi = c(0,0,0,0))
plot(s_dat[2,], r_dat[2,], xlim = c(0,xM), ylim = c(0,yM), type = "n",
     xaxs = "i", yaxs = "i", cex.lab = 1.2,
     xlab = expression(Spawners~(10^3)),
     ylab = expression(Recruits~(10^3)),
     xaxt = "n", yaxt = "n", bty="L")
axis(1, at = seq(0,xM,dd*2), labels = seq(0,xM,dd*2)/1000)
axis(2, at = seq(0,yM,dd*2), labels = seq(0,yM,dd*2)/1000, las=1)
for(i in 1:length(aa)) {
```

```r
  lines(exp(aa[i]) * seq(0,xM) / (1 + bb * seq(0,xM)),
        col = "darkgray")
}
abline(a = 0,b = 1,lty = "dashed")

## add S-R estimates and medians
nCB <- n_yrs-age_max
## years with complete returns
points(s_dat[2, 1:nCB], r_dat[2, 1:nCB],
       xlim = c(0,xM), ylim = c(0,yM),
       pch = 16, col = "blue3")
segments(s_dat[2, 1:nCB], r_dat[1, 1:nCB],
         s_dat[2, 1:nCB], r_dat[3, 1:nCB],
         col = "blue3")
segments(s_dat[1, 1:nCB], r_dat[2, 1:nCB],
         s_dat[3, 1:nCB], r_dat[2, 1:nCB],
         col = "blue3")
nTB <- dim(s_dat)[2]
## years with incomplete returns
segments(s_dat[2, (nCB+1):nTB], r_dat[1, (nCB+1):nTB],
         s_dat[2, (nCB+1):nTB], r_dat[3, (nCB+1):nTB],
         col = clr)
segments(s_dat[1, (nCB+1):nTB], r_dat[2, (nCB+1):nTB],
         s_dat[3, (nCB+1):nTB], r_dat[2, (nCB+1):nTB],
         col = clr)
points(s_dat[2, (nCB+1):nTB],r_dat[2, (nCB+1):nTB],
       xlim = c(0,xM), ylim = c(0,yM),
       pch = 16, col = clr)
text(x = par()$usr[1] + diff(par()$usr[1:2]) * xoffSet,
     y = par()$usr[4] - diff(par()$usr[3:4]) * yoffSet,
     "(a)")

## posterior for alpha
clr <- rgb(0, 0, 255, alpha = 50, maxColorValue = 255)
a_thresh <- 59
par(mai = c(0.8,0.4,0.3,0.1))
## B-H alpha
R_alpha_est <- mod_res[, "alpha"]
alphaCI <- quantile(R_alpha_est, CI_vec)
R_alpha_est[R_alpha_est > a_thresh] <- a_thresh
hist(R_alpha_est, freq = FALSE, breaks = seq(0, a_thresh+1, 2),
     col = clr, border = "blue3",
     xlab = "", ylab = "", main = "", cex.lab = 1.2, yaxt = "n")
aHt <- (par()$usr[4]-par()$usr[3])/12
arrows(alphaCI, par()$usr[3], alphaCI,par()$usr[3]-aHt,
       code = 1, length = 0.05, xpd = NA, col = "blue3", lwd = 1.5)
mtext(expression(Instrinsic~productivity~(alpha)), 1, line = 3, cex = 1)
text(x = par()$usr[1],
```

```
     y = par()$usr[4] * 1.05,
     "(b)", xpd=NA)

## posterior for K
par(mai = c(0.8,0.4,0.3,0.1))
aa <- mod_res[, "alpha"]
bb <- mod_res[, "beta"]
## K in 1000s
R_b_est <- (aa-1) / bb / 1000
R_b_est <- R_b_est[R_b_est > 0]
R_b_CI <- quantile(R_b_est, CI_vec)
## pile into last ban for plotting
R_b_est[R_b_est > 13] <- 13
brks <- seq(around(min(R_b_est), "floor"),
            around(max(R_b_est), "ceiling"),
            length.out = length(seq(0, a_thresh, 2)))
hist(R_b_est, freq = FALSE, breaks = brks, col = clr, border = "blue3",
     xlab = "", xaxt = "n", yaxt = "n",
     main = "", ylab = "", cex.lab = 1.2)
axis(1, at = seq(around(min(R_b_est), "floor"),
                 around(max(R_b_est), "ceiling"),
                 2))
aHt <- (par()$usr[4] - par()$usr[3]) / 12
arrows(R_b_CI, par()$usr[3], R_b_CI,par()$usr[3]-aHt,
       code = 1, length = 0.05, xpd = NA, col = "blue3", lwd = 1.5)
mtext(expression(paste("Carrying capacity (",italic(K),", ",10^3,")")),
      side = 1, line = 3, cex = 1)
text(x = par()$usr[1],
     y = par()$usr[4] * 1.05,
     "(c)", xpd=NA)
```

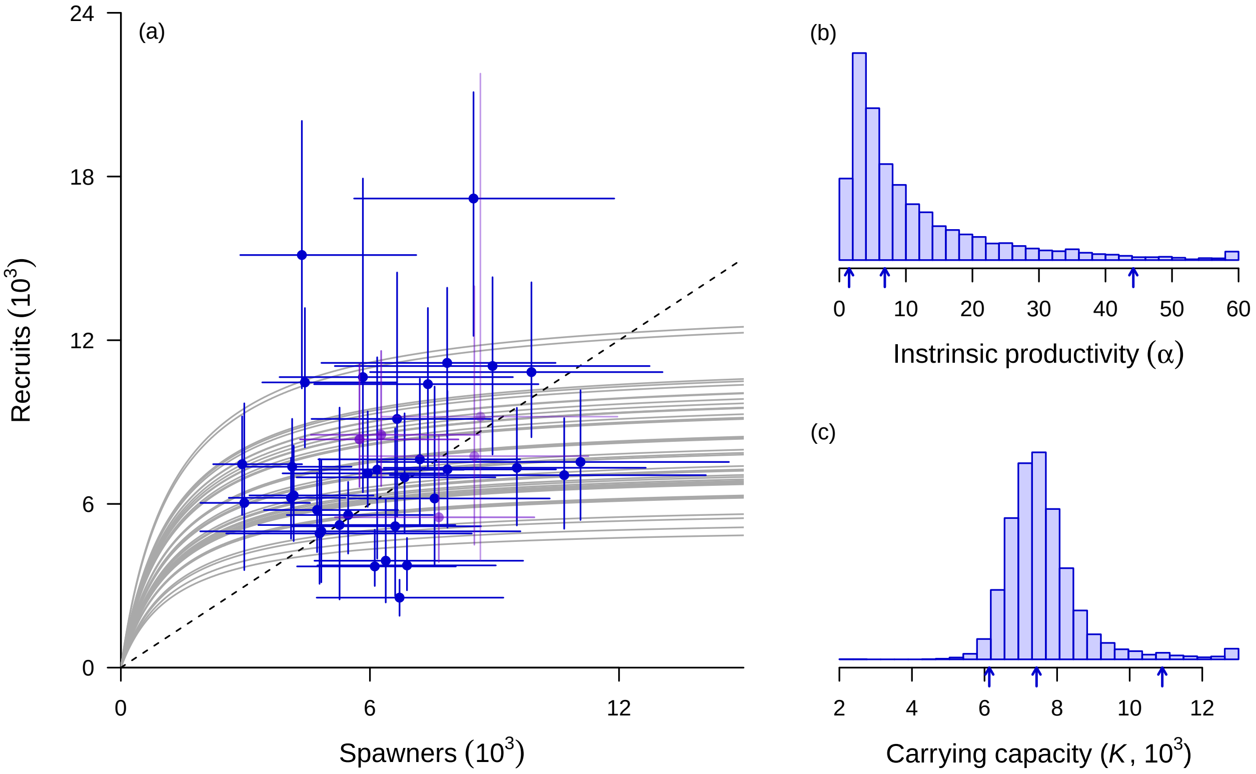

Figure 3. Relationship between the number of spawning adults and their subsequent surviving offspring (recruits), assuming mean values for all covariates (a); and the estimated posterior distributions for the intrinsic productivity (b) and carrying capacity (c). Points in (a) are medians of the posterior estimates; error bars indicate the 95% credible intervals. Blue points are for estimates with complete broods; purple points are for the most recent years with incomplete broods. Gray lines show the median relationship for each of the 41 years in the time series based on annual model estimates of productivity. Note that for plotting purposes only in (b) and (c), the density in the largest bin for each parameter contains counts for all values greater than or equal to it. Vertical arrows under the x-axes in (b) and (c) indicate the $2.5^{\text{th}}$, $50^{\text{th}}$, and $97.5^{\text{th}}$ percentiles.

Here are summaries of the posterior distributions for $\alpha$ and $K$.

```
## intrinsic productivity
round(alphaCI, 2)
```

```
##  2.5%   50% 97.5%
##  1.46  6.83 44.18
```

```
## carrying capacity
round(R_b_CI, 2)
```

```
##  2.5%   50% 97.5%
##  6.13  7.43 10.90
```

##### 4.3 Covariate effects

Here are time series plots of the covariates (a-c) and histograms of their effects on productivity (d-f).

```r
clr <- rgb(0, 0, 255, alpha = 50, maxColorValue = 255)
xoffSet <- 0.04
yoffSet <- 0.03

par(mfrow=c(n_cov,2), mai=c(0.4,0.2,0.1,0.1), omi=c(0.2,0.5,0,0))

c_est <- mod_res[,grep("gamma", colnames(mod_res))]
ylN <- floor(min(c_est)*10)/10
ylM <- ceiling(max(c_est)*10)/10
brks <- seq(ylN,ylM,length.out=diff(c(ylN,ylM))*40+1)
t_idx <- seq(yr_frst,length.out=n_yrs-age_min+n_fore)
dat_cvrs <- as.matrix(dat_cvrs[seq(length(t_idx)),])

for(i in 1:n_cov) {
  if(i==4) {
    dat_cvrs[,i+1] <- dat_cvrs[,i+1]/1000
  }
  ## plot covar ts
  plot(dat_cvrs[, "year"], dat_cvrs[, i+1],
       pch = 16, col = "blue3", type = "o",
       xlab = "", ylab = "", main = "", bty = "L",
       cex.axis = 1.2)
  text(x = par()$usr[1] + diff(par()$usr[1:2]) * xoffSet,
       y = par()$usr[4] - diff(par()$usr[3:4]) * yoffSet,
       paste0("(",letters[i],")"),
       cex = 1.2)
  mtext(side = 2, cov_names[i], line = 3, cex = 1.2)
  if(i == n_cov) {
    mtext(side = 1, "Brood year", line = 3)
  }
  ## plot covar effect
  hist(c_est[,i],
       freq = FALSE, breaks = brks, col = clr, border =" blue3",
       xlab = "", yaxt = "n", main = "", ylab = "", cex.axis = 1.2)
  c_CI <- quantile(c_est[,i],CI_vec)
  aHt <- (par()$usr[4]-par()$usr[3])/20
  arrows(c_CI, par()$usr[3]-0.005, c_CI, par()$usr[3] - aHt,
         code = 1,length = 0.05, xpd = NA, col = "blue3", lwd = 1.5)
  abline(v = 0, lty = "dashed")
  text(x = par()$usr[1] + diff(par()$usr[1:2]) * xoffSet,
       y = par()$usr[4] - diff(par()$usr[3:4]) * yoffSet,
       paste0("(",letters[i+n_cov],")"),
       cex = 1.2)
  if(i == n_cov) { mtext(side = 1,"Effect size", line = 3) }
}
```

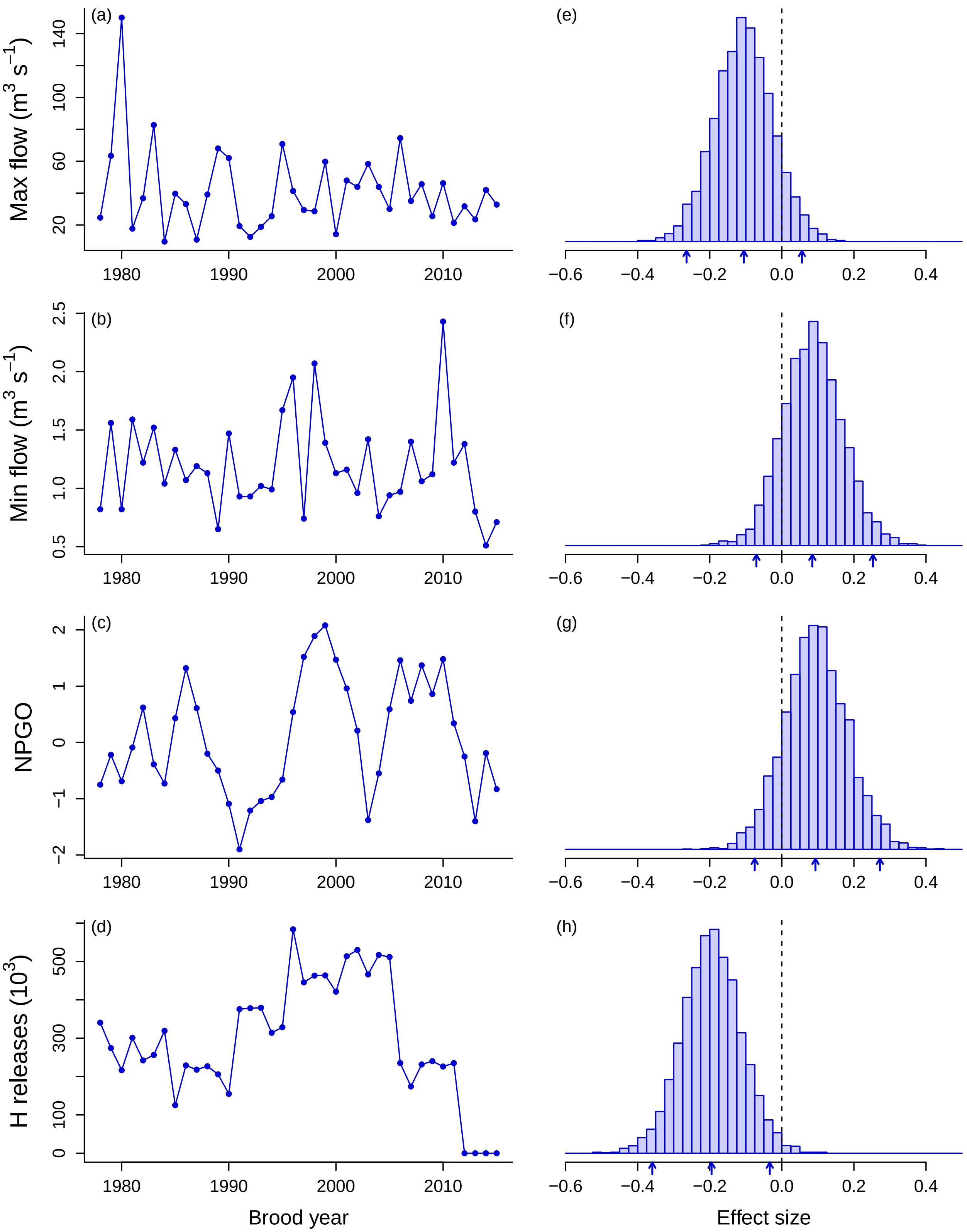

Figure 4. Time series of the environmental covariates used in the model (a-d), and their estimated effects on population productivity (e-g). Small arrows under histograms denote 2.5$^{\text{th}}$, 50$^{\text{th}}$, and 97.5$^{\text{th}}$ percentiles of the posterior distribution.

Here is a summary of the covariate effect sizes

```
gamma_CI <- apply(c_est, 2, quantile, c(2.5, 5, 50, 95, 97.5)/100)
t(round(gamma_CI, 2))
```

```
##           2.5%    5%   50%   95% 97.5%
## gamma[1] -0.26 -0.24 -0.11  0.03  0.06
## gamma[2] -0.07 -0.05  0.08  0.22  0.25
## gamma[3] -0.08 -0.05  0.09  0.24  0.27
## gamma[4] -0.36 -0.33 -0.20 -0.06 -0.03
```

##### 4.4 Process errors

Here is the time series of the residuals from the process model. They represent the population's productivity after accounting for the effects of density dependence and environmental covariates.

```
## time sequence
t_idx_a <- seq(yr_frst, length.out = n_yrs-age_min+n_fore)
## plot data
p_dat <- mod_res[, grep("res_ln_Rec", colnames(mod_res))]
p_dat <- apply(p_dat, 2, quantile, CI_vec)
yp_min <- min(p_dat)
yp_max <- max(p_dat)
## plot
par(mai = c(0.8,0.8,0.1,0.1), omi = c(0,0.2,0.1,0.2))
plot(t_idx_a, p_dat[3,],
     type = "n",  bty = "L",
     ylim = c(yp_min,yp_max),
     xlab = "Brood year", ylab = "Process error", main = "",
     cex.lab = 1.2)
abline(h = 0, lty = "dashed")
polygon(c(t_idx_a, rev(t_idx_a)), c(p_dat[3,], rev(p_dat[1,])),
        col = clr, border = NA)
lines(t_idx_a, p_dat[2,], col = "blue3", lwd = 2)
```

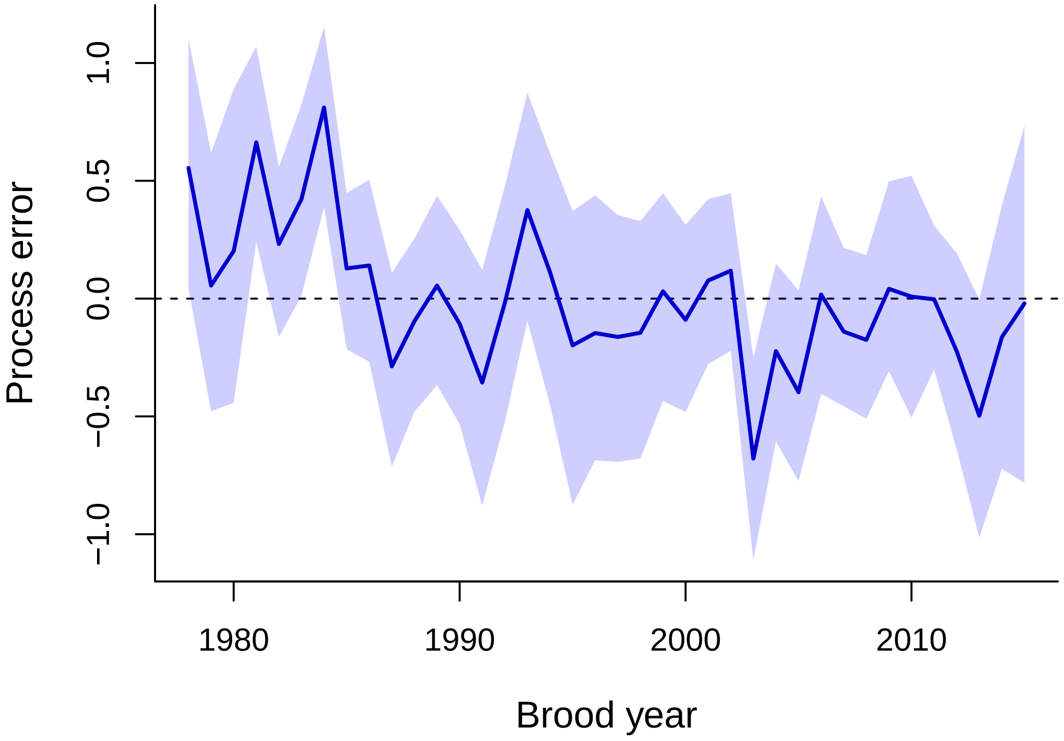

Figure 5. Time series of the estimated process errors, which represent the population's productivity after accounting for the effects of density dependence and environmental covariates. The solid line is the median estimate and the shaded region indicates the 95% credible interval.

##### 4.5 Management reference points

Here are a number of management reference points.

```
## abbreviations for ref points
ref_names <- c("MSY", "Smsy", "Umsy", "Umax")
## proportions of MSY to consider
yld_prop <- c(0.75, 0.85, 0.95)
## median values for a & b
aa <- mod_res[, grep("E_BH_a", colnames(mod_res))]
alpha <- exp(aa)
mcmc <- length(aa)
beta <- mod_res[, grep("beta", colnames(mod_res))]

## empty matrix for ref pts
ref_pts <- matrix(NA, mcmc, length(ref_names))
colnames(ref_pts) <- ref_names
## spawner series for optimal yield profile
SS <- seq(100, 1e4, 100)
## empty matrix for optimal yield profiles
OYP <- matrix(0, length(SS), length(yld_prop))
for(i in 1:mcmc) {
```

```
    ## spawners at MSY
    ref_pts[i, "Smsy"] <- (alpha[i] / beta[i]) * sqrt(1 / alpha[i]) - (1 / beta[i])
    ## MSY
    ref_pts[i, "MSY"] <- (ref_pts[i,"Smsy"] * alpha[i]) /
                         (1 + beta[i] * ref_pts[i, "Smsy"]) - ref_pts[i, "Smsy"]
    ## harvest rate at MSY
    ref_pts[i, "Umsy"] <- 1 - sqrt(1 / alpha[i])
    ## max harvest rate
    ref_pts[i, "Umax"] <- 1 - 1/alpha[i]
    ## yield over varying S
    yield <- ((SS * alpha[i]) / (1 + beta[i] * SS)) - SS
    for(j in 1:length(yld_prop)) {
        OYP[,j] <- OYP[,j] + 1*(yield > yld_prop[j] * ref_pts[i, "MSY"])
    }
}
OYP <- OYP/mcmc

## Prob of overfishing
hh <- seq(100)
Pr_over <- cbind(hh,hh,hh)
colnames(Pr_over) <- c("Umsy75","Umsy","Umax")
for(i in hh) {
  Pr_over[i,"Umsy75"] <- sum(ref_pts[,"Umsy"] * 0.75 < i/100)/mcmc
  Pr_over[i,"Umsy"] <- sum(ref_pts[,"Umsy"] < i/100)/mcmc
  Pr_over[i,"Umax"] <- sum(ref_pts[,"Umax"] < i/100)/mcmc
}

## posterior exploitation rate & spawner abundance
aer <- Sp_ts <- mod_res[,grep("Sp", colnames(mod_res))]
for(i in 1:n_yrs) {
    aer[,i] <- dat_harv[i] / (dat_harv[i] + Sp_ts[,i])
}

layout(matrix(c(2, 1, 4, 3), 2, 2), heights = c(1, 5))
yoffSet <- 0.10
yoffSet <- 0.05

## (a) Optimal yield profile
par(mai=c(0.9, 0.9, 0, 0), omi=c(0, 0, 0.1, 0.1))
x_lp <- yld_prop
for(i in 1:length(x_lp)) {
    x_lp[i] <- SS[max(which(OYP[,i] == max(OYP[,i])
                        | abs(OYP[,i] - (yld_prop[i]-0.3)) <= 0.05))]
}
matplot(SS, OYP, type="l", lty="solid",  ylim=c(0,1),
        col=c("slateblue","blue","darkblue"), lwd=2,
        xlab = "Spawners", ylab = "Probability of X% of MSY", main = "",
        las=1, cex.lab=1.2)
```

```
points(x = x_lp, y = yld_prop-0.3,
       pch = 21, cex = 3.5,
       col = "white", bg = "white")
text(x = x_lp, y = yld_prop-0.3, paste0(yld_prop*100, "%"),
     col=c("slateblue","blue","darkblue"), cex=0.7)
text(x = par()$usr[1] + xoffSet * diff(par()$usr[1:2]),
     y = par()$usr[4] - yoffSet * diff(par()$usr[3:4]),
     "(a)")
## marginal histogram of posterior spawner abundances
par(mai=c(0, 0.9, 0.05, 0))
hist(Sp_ts[Sp_ts<1e4], breaks = 40,
     col = clr, border = "blue3",
     yaxs = "i", xaxt = "n", yaxt = "n",
     main = "", ylab = "")

## (b) Probability of overfishing
par(mai=c(0.9, 0.9, 0, 0))
matplot(Pr_over, type = "l", lwd = 2, lty = "solid",
        col = c("slateblue","blue","darkblue"),
        ylab="Probability of overfishing",
        xlab="Harvest rate", xaxt="n",
        las = 1, cex.lab = 1.2)
axis(1, seq(0,100,20), seq(0,100,20)/100)
x_lp <- c(0, 0, 0)
for(i in 1:length(x_lp)) {
  x_lp[i] <- max(which(abs(Pr_over[,i] - 0.5) <= 0.05))
}
points(x = x_lp, y = rep(0.5, 3), pch = 21, cex = 4,
       col = "white", bg = "white")
text(x = x_lp, y = 0.5, expression(U[M75], U[MSY], U[Max]),
     col = c("slateblue", "blue", "darkblue"), cex = 0.8)
text(x = par()$usr[1] + xoffSet * diff(par()$usr[1:2]),
     y = par()$usr[4] - yoffSet * diff(par()$usr[3:4]),
     "(b)")
## marginal histogram of posterior harvest rates
par(mai = c(0 ,0.9, 0.05, 0))
hist(aer, breaks = seq(0, 40)/40,
     col = clr, border = "blue3",
     yaxs = "i", xaxt = "n", yaxt = "n",
     main = "", ylab = "")
```

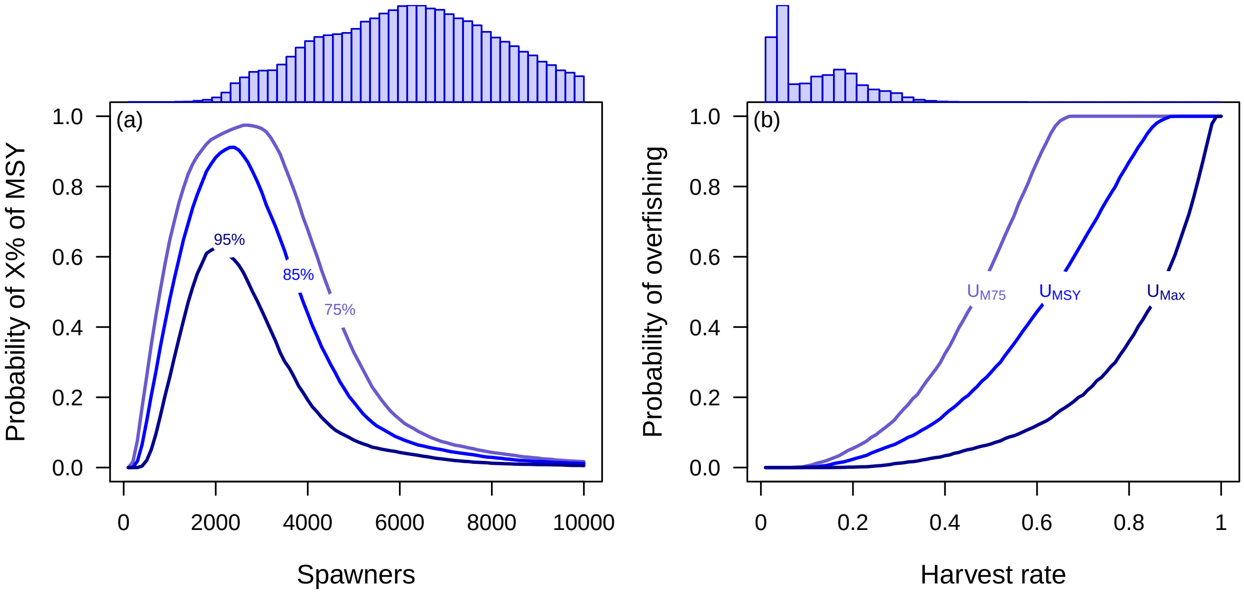

Figure 6. Plots of (a) the probability that a given number of spawners produces average yields achieving 95%, 85%, or 75% of the estimated maximum sustainable yield (MSY); and (b) the cumulative probability of overfishing the population, based on harvest rates equal to those at 75% of MSY, at MSY, and at the maximum per recruit. The histograms above (a) and (b) are distributions of the posterior estimates for the number of spawners and harvest rates, respectively; the histogram in (a) has been truncated at $10^4$.
